# Population dynamics in spatial suppression gene drive models and the effect of resistance, density dependence, and life history

**DOI:** 10.1101/2024.08.14.607913

**Authors:** Xinyue Zhang, Weitang Sun, Isabel K. Kim, Philipp W. Messer, Jackson Champer

## Abstract

Due to their super-Mendelian inheritance, gene drive systems have the potential to provide revolutionary solutions to critical public health and environmental problems. For suppression drives, however, spatial structure can cause “chasing” population dynamics that may postpone target population elimination or even cause the drive to fail. In chasing, wild-type individuals elude the drive and recolonize previously suppressed areas. The drive can re-enter these recolonized areas, but often is not able to catch up to wild-type and finally eliminate it. Previous methods for chasing detection are only suitable to limited parameter ranges. In this study with expanded parameter ranges, we found that the shift from chasing dynamics to static equilibrium outcomes is continuous as drive performance is reduced. To quantify this, we defined a Weighted Average Nearest Neighbor statistic to assess the clustering degree during chasing, while also characterizing chasing by the per-generation chance of population elimination and drive loss. To detect chasing dynamics in local areas and to detect the start of chasing, we implemented Density-Based Spatial Clustering of Applications with Noise. Using these techniques, we determined the effect of arena size, resistance allele formation rate in both the germline and in the early embryo from maternally deposited Cas9, life history and reproduction strategies, and density-dependent growth curve shape on chasing outcomes. We found that larger real-world areas will be much more vulnerable to chasing and that species with overlapping generations, fecundity-based density dependence, and concave density-dependent growth curves have smaller and more clustered local chasing with a greater chance of eventual population elimination. We also found that embryo resistance and germline resistance hinder drive performance in different ways. These considerations will be important for determining the necessary drive performance parameters needed for success in different species, and whether future drives could potentially be considered as release candidates.

## 1 Introduction

Gene drive systems have been rapidly developed due to their enormous potential for dealing with issues from disease vectors and other pests (Gould 2008; Champer *et al*. 2016). Gene drive systems use synthetic genetic elements designed to increase in frequency over time in a population, spreading a desired allele via non-Mendelian inheritance (DiCarlo *et al*. 2015). CRISPR gene editing technology has greatly facilitated development of engineered gene drive (Bier 2021). For example, a modification drive could be designed to spread a gene through a mosquito population that prevents disease transmission (Carballar-Lejarazú & James 2017). A powerful suppression drive can potentially eliminate entire populations of disease vectors, pest species, and invasive species (Godwin *et al*. 2019; Champer *et al*. 2021a; Ferreira-Martins *et al*. 2021). A homing suppression drive targeting an essential female fertility gene has successfully eliminated cage populations of *Anopheles gambiae* mosquitoes (Kyrou *et al*. 2018). However, the ethically-responsible development and deployment of gene drives limits field experiments (Kc *et al*. 2020; Kormos *et al*. 2022), making computational models an important approach to anticipate their performance in a natural environment.

Spatial population structure has been shown to have a major impact on the outcome of suppression drive in several models (North *et al*. 2019; Champer *et al*. 2021a; Girardin & Débarre 2021; Paril & Phillips 2022). Drive systems that are capable of eliminating a panmictic population can fail due to spatial structure (Paril & Phillips 2022). Specifically, partial population suppression can lead to an unstable coexistence of drive and wild-type alleles due to the limited dispersal distance (Champer *et al*. 2021a; Birand *et al*. 2022). Because dispersal is constrained by geography and the movement of individuals, the spread of drive alleles is also constrained, allowing wild-type individuals to escape into empty areas that have previously been suppressed by the drive. The drive then has to “chase” the wild-type allele over the landscape. When analyzing the dynamics of failed suppression drives, chasing dynamics appeared to be a major cause of failure for efficient drives in spatial models (Bull *et al*. 2019; Champer *et al*. 2021a; Olejarz & Nowak 2024).

To construct successful suppression drives for practical applications, it is important to have a better understanding of chasing dynamics and what factors influence them. Chasing is generally considered different from a static equilibrium by its large variance in population density over time and space (Champer *et al*. 2021a; Kim *et al*. 2023). An equilibrium outcome involves reduced population, but no local elimination except potentially from stochastic and short-term effects. A previous study of suppression drive in continuous space developed an ad-hoc procedure to detect if and when chasing occurred using Green’s coefficient as a measure of population clustering and wild-type population size (Champer *et al*. 2021a). However, it was specific to the parameter space in the study and not tested over the full range of possible outcomes and models.

Other studies have covered the importance of ecological factors and resistance in suppression drive performance, such as density-dependent factors and the specific negative effects of density, species interactions, spatial structure within and between populations, and mating behavior (Godfray *et al*. 2017; Dhole *et al*. 2020). However, these ecological factors have not been studied in the context of chasing dynamics. Density dependence is an important factor for regulating populations and is particularly important in suppression drives where the density can rapidly change. For example, the Allee effect reduces population growth rate at very low population density (Stephens *et al*. 1999; Barton & Turelli 2011). In spatial models, drive loss is more likely to occur when Allee effects are strong due to lack of ability to find mates (North *et al*. 2019; Champer *et al*. 2021a; Birand *et al*. 2022). The suppression performance of a similar drive on two populations with different density-dependent curves may have considerable differences (Dhole *et al*. 2020). The lifecycle phase where density-dependence acts can also matter. Some species may decrease their number of offspring to continue effectively reproducing when adults experience high competition. Others can produce many offspring, yet only a small portion of them typically survive at high density due to direct resource competition between juveniles (Roff 1993).

Furthermore, there are other aspects of drive performance and ecology that have not been thoroughly investigated because of computational resource limits (Champer *et al*. 2022). Larger spatial areas, for example, may greatly facilitate chasing. Nonfunctional resistance alleles caused by end-joining repair in the germline or in embryos from maternally deposited Cas9 could also impede the spread of the drive (Beaghton *et al*. 2019; Champer *et al*. 2021a) and have been experimentally found over a fairly wide range (Champer *et al*. 2019; Hammond *et al*. 2021; Du *et al*. 2024).

In this study, we sought to better characterize chasing and spatial suppression drive outcomes in general. We defined a continuous variable using the definition of average nearest neighbor ratio to quantify chasing dynamics (Chester 1987), which describes the clustering degree of the population distribution. We found that the change from chasing dynamics to widespread equilibrium outcome from less efficient drives is continuous, with slightly subpar drives (that lack the ability to eliminate panmictic populations) often having continuous local extinction/recolonization cycles from stochastic effects. We also developed a general method to detect whether a given local area is experiencing chasing. This method can allow for a more sophisticated understanding of both the general and the local distribution of the chasing dynamics. Using these assessment and detection methods, we investigate how chasing dynamics are impacted by ecology, density-dependence, and previously unconsidered drive performance parameters. We specifically examine arena size, resistance allele formation rate, life history, reproduction strategies, and density-dependent growth curve shape. We devote particular attention to explaining why a previous mosquito-specific model showed higher rates of chasing than a discrete-generation model (Champer *et al*. 2022).

## 2 Methods

### 2.1 Population model

In this study, we use an individual-based population model simulating a sexually reproducing diploid population. The model is general and can potentially be applied to a variety of species. Time steps could represent discrete units of time with overlapping generations, or they could represent discrete generations, with only newly produced offspring surviving to the next time step. Competition in the model affects either fecundity or offspring viability to simulate different reproduction strategies. We use a two-dimensional continuous space model where every individual’s position is recorded. Initially, wild-type individuals are randomly distributed in the square landscape of length 1. Females choose their mates randomly within a radius specified by the mating distance, which is equal to the average dispersal distance parameter and is set to 0.05. If females cannot find a mate, they will not reproduce in this time step. The number of offspring is dependent on the reproduction strategy (see below). The positions of offspring are offset from their mother with a distance drawn from a normal distribution with an average equal to the *dispersion distance* in each dimension. Surviving adults migrate the same way in each time step. If any individual is placed outside the area, its position is redrawn until it falls within the arena.

In the fecundity model, total carrying capacity *K* is set to 20,000. Females generate a number of offspring based on their fitness and local competition. In our spatial model, the competition is limited in radius specified by the interaction distance parameter, which is set to 0.01. The local density of each position *ρ*_*i*_ is defined as the density of individuals within a circle of radius equal to interaction distance around it. The expected average density of individuals at normal equilibrium is *ρ*_*k*_ = *K*/*area*. The relative reproduction of the female is then calculated by:

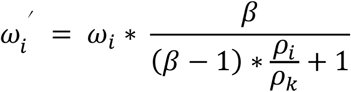

where *β* represents the low-density growth rate of the population (normally set to 6). The actual number of offspring is drawn from a binomial distribution with a number of draws equal to *β*_*max*_. The chance of each draw producing an offspring is 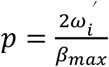, where *β*_*max*_represents the maximum number of offspring (set to 25 by default), *ω*_*i*_ represents the genotype fitness of female. This ensures that every female produces an average of two offspring per generation under normal conditions (no fitness costs, population density equal to the carrying density).

In contrast, individuals produce many offspring in the viability model. The number of offspring is drawn from a Poisson distribution with and average *λ* = 2 * *β*_*max*_ * *ω*_*i*_. After reproduction, offspring compete to survive, and the survival rate is:

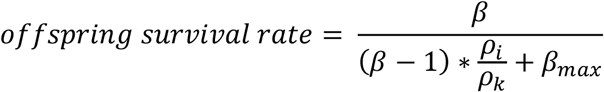

where *ρ*_*i*_ is defined as the density of newborn offspring within a circle of radius equal to interaction distance around it. Offspring become adults starting at the next time step.

In the discrete-generation model, the generations are non-overlapping. Every individual can only survive and reproduce in one time, and the offspring become the reproducing adults in the next time step. However, many species continuously reproduce and have overlapping generations. To create an overlapping-generations model, the survival rates of individuals are age-based, the survival rates at each age are given as follows:

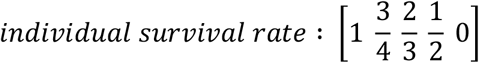

This results in a linear decrease through time in the number of individuals in a single age class. Individuals can mate and reproduce in every time step, except for the time step in which they are born. In the fecundity model, the number of eggs during reproduction is drawn from a binomial distribution with a number of draws equal to *β*_*max*_/*generation*_*time*. The chance of each draw producing an offspring is 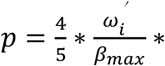 in the overlapping-generations fecundity model. In viability model, the number of offspring is drawn from a Poisson distribution with an average of 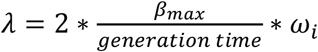, and the survival rate of offspring in the overlapping-generations viability model is equal to 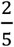 multiplied by the offspring survival rate in the discrete-generation viability model.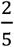 represents the proportion of adults that are age 1 out of all adults when the population is at equilibrium. Individuals migrate with average dispersal distance 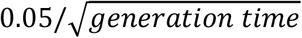, which produced the same average per-generation dispersal as the discrete generation model. The generation time of this overlapping model is two time steps. We run the overlapping-generation model for 750 time steps, and we run the discrete generation model for 375 generations. Simulation can end early if the population is eliminated, or if the drive is lost. We recorded frequencies of different alleles, population size, the number of fertile females, and the average nearest neighbor ratio for each time step. See **Table S1** for a list of default parameters

### 2.2 Density-dependent growth curves

The density-dependent growth curve describes the relationship between population size and population growth rate. Higher growth at lower population density can contribute to chasing. Population growth rate is defined as *N*_*t*_/*N*_*t*-1_, where *N*_*t*_ represents the population size of generation *t*. We investigated the influence of three density-dependent growth curves of different shapes to the outcomes of suppression drive in the discrete-generation fecundity model. Our default Beverton-Holt growth curve is denoted above (Beverton & Holt 1957). A general *θ* −logistic model is written as *dN*⁄*dt* = *rN*(1 − (*N*⁄*K*)^*θ*^), where *r* is the growth rate, K is the carrying capacity, and *θ* is a parameter that determines the growth curve (Gilpin & Ayala 1973). When *θ* = 1, the growth rate linearly declines as the population size increases (Cook 1965). Connecting Ricker’s model (Ricker 1954; Cook 1965) and studies of laboratory *Drosophila* populations (Gilpin & Ayala 1973), the *θ*-Ricker model (Thomas *et al*. 1980) was put forward as:

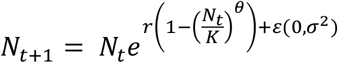

The parameter *θ* determines the shape of the density-dependent growth curve, reflecting the degree of influence to population size as abundance interacts with resource availability (Freckleton *et al*. 2003) and type of competition within species (Johst *et al*. 2008). When *θ* < 1, the growth rate declines sharply when the population size is low, which makes it harder to recover from low population density. Some laboratory populations show concave density-dependent growth curves (Hassell *et al*. 1976), similar to our Beverton-Holt growth curve. When *θ* > 1, the growth rate remains at a high level for longer when the population size is low, which can make the population harder to eliminate. The density-dependent growth curves of some large mammalian herbivores are convex (Owen-Smith 2006).

Here, we changed the reproductive rate of females to represent different density-dependent growth curves. The default 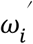 indicated above results in a concave density-dependent growth curve. The linear and convex density-dependent growth curves are set as:

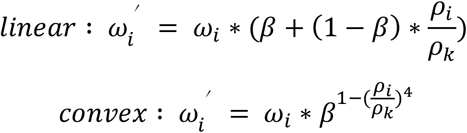

### 2.3 Suppression drive strategies

Female fertility homing drive can decrease the number of offspring by making females sterile. It is also the best studied drive from an experimental perspective (Kyrou *et al*. 2018; Yang *et al*. 2022). In a previous modeling study, such a drive performed well, but was still not powerful enough to eliminate the population for much of the parameter space (Champer *et al*. 2021a; Champer *et al*. 2022).

This CRISPR/Cas9 homing drive targets a haplosufficient but essential female fertility gene. Females are sterile if they lack at least one wild-type or functional resistance allele. In drive heterozygotes, cleavage and homology-directed repair happen in the germline, which will convert wild-type alleles into drive alleles. If homology-directed repair fails, end-joining will cause the guide RNA (gRNA) site to be mutated and create resistance alleles that cannot be cleaved. Resistance alleles mostly are nonfunctional, called “r2” resistance alleles. Some resistance alleles called “r1” can maintain target gene function. “r1” resistance alleles play a decisive role in preventing population suppression. If they form at even a low probability, the population is likely to recover (Birand *et al*. 2022). In our simulations, we neglect r1 alleles because they can be avoided in homing suppression drives by using multiplexed gRNAs (Prowse *et al*. 2017; Bishop *et al*. 2022; Yang *et al*. 2022) and conserved target sites (Kyrou *et al*. 2018).

In our model, wild-type alleles have a probability equal to the *drive conversion* parameter to be converted into a drive allele in heterozygotes. Half of the remaining wild-type alleles are converted to r2 resistance alleles by default (Hammond *et al*. 2021). However, this can also be a variable parameter, and the *germline resistance* rate is the percentage of wild-type alleles converted into r2 resistance alleles. Together, the *drive conversion* and *germline resistance* rates cannot exceed 1. If the mother has a drive allele, offspring may have deposited Cas9 and gRNA, which converts wild-type alleles to r2 resistance alleles at a probability equal to the *embryo resistance* allele formation rate. Because somatic or other undesired Cas9 cleavage can reduce the reproduction rate of drive females, we set a variable drive *female heterozygote fitness* to be less than 1 (which is wild-type fitness) when modeling this phenomenon.

The initial release number of female fertility homing drive does not influence the final suppression level and outcome in most situations (North *et al*. 2020). We thus use a simple release of drive heterozygotes at a frequency of 10 percent of the carrying capacity, all within a central circle of radius 0.01.

### 2.4 Local chasing detection

To determine where and when chasing occurs, we divided the space into squares with lengths equal to the *average dispersal distance*. Together with the density interaction distance, this sets the scale for the simulation and is specifically the smallest distance that should have a significant effect on movement of individuals. A key feature of chasing is that wild-type individuals escape from drive-populated areas into empty areas previously cleared by the drive (Champer *et al*. 2021a). Thus, we define chasing as occurring in a cell if it is first eliminated (the number of individuals in this cell is zero) and then recolonized such that the number of non-drive individuals is larger than the average number of non-drive individuals across all cells. In other words, we define chasing to be when wild-type individuals reoccupy one cell and reach a higher density of non-drive individuals than the average level of the whole space. However, we also need to ensure that we don’t flag short-term dynamics when the chasing equilibrium may still be developing as chasing (in this situation, the population may still be rapidly eliminated before a chasing equilibrium is established across the whole arena). Thus, we also require that the number of wild-type individuals is larger than 10 percent of the population size. We further require that at least 4 cells are flagged as chasing to consider the whole system in a state of chasing.

The positions of detected chasing cells can show the directions of wild-type escape from the center of the wild-type clusters to the positions of the chasing cells (**Figure 1A**).

**Figure 1.**
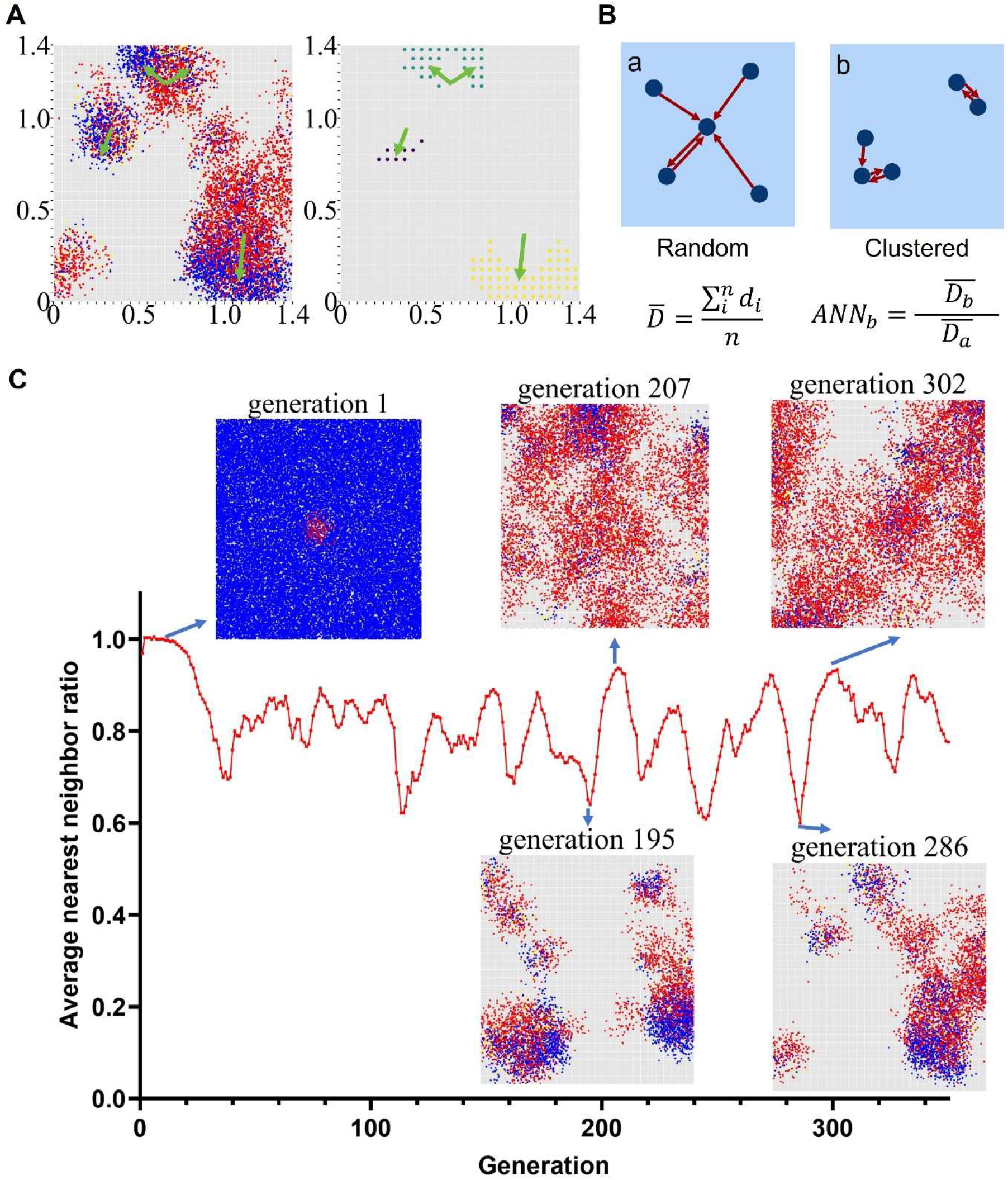
The average nearest neighbor ratio and the chasing clustering method. (**A**) An example of chasing. The left graph shows the distribution of all individuals across the whole area. Blue dots represent wild-type individuals, red dots represent drive carriers, and yellow dots represent individuals with r2 alleles. Green arrows show the escape directions of the wild-type individuals. The right graph shows chasing cells across the area detected by the local chasing detection algorithm in. Different colors represent different local chasing clusters. (**B**) The calculation of average nearest neighbor ratio. The blue dots represent the five individuals, and the red arrows point to their nearest neighbors. (**C**) The average nearest neighbor ratio during chasing and several corresponding snapshots of the population at peaks and troughs of the *ANN* ratio.

### 2.5 Drive spread speed measurements

To evaluate the wave speed that wild-type individuals spread into the empty space, we divided a one-dimensional space into slices with equal length and then released wild-type individuals evenly in the leftmost slices. We used the number of generations that wild-type individuals take to reach the rightmost slice of the area to evaluate the wave spread speed.

Chasing dynamics not only includes the process in which wild-type individuals spread into empty areas but also includes the process in which the drive spreads into areas with wild-type individuals. To evaluate the change of drive alleles and nonfunctional resistance alleles in the two-dimensional population model, the relative increase rate of allele frequency of each generation is defined as (*D*_*t*+1_ − *D*_*t*_)/*D*_*t*_, where *D*_*t*_ represents the specific allele frequency of generation *t*. In both the panmictic model and spatial models, we found that the relative increase rate of drive allele frequency linearly declines with increasing drive allele frequency, and r2 resistance alleles also follow this rule (Figure S2). Thus, we used a linear fit of the data when the drive allele frequency was between 0.05 and 0.9 using Huber loss function and Nelder-Mead simplex algorithm (Wright 1996) to optimize it.

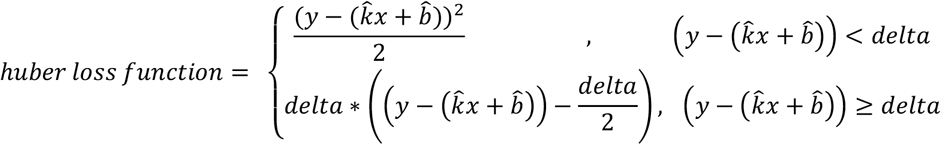

where delta is 0.005. In the initial generations after the release, because most individuals with drive alleles are heterozygous, the drive allele spreads at a higher rate than in other situations where the population reaches a balance that includes drive homozygotes and resistance alleles. Additionally, when the frequency of the drive allele is high, there are more data points, which is likely to influence the fitting results. To address this, we separated all data into 8 even ranges based on the drive allele frequency and sampled the same number within each range, except for the first and last range (representing drives that were artificially in a heterozygote-only starting situation and an equilibrium near the end with small sample sizes). Finally, we recorded the zero-frequency relative increase rate (vertical intercept of the linear relationship) and maximum drive frequency (horizontal intercept of the linear relationship). The zero-frequency relative increase rate represents the drive spread speed when the drive frequency is low. The maximum drive frequency represents the drive frequency when the population reaches equilibrium (assuming that the population persists and is not eliminated), and it is somewhat representative of the long-term power of the drive to eliminate a population. If more than 70% of generations have a drive allele frequency of less than 0.1 (which did not occur in the parameter range studied in this manuscript), the relative increase rate of the drive allele appears to be distributed randomly when the drive frequency is low. In this case, we increased the release frequency from 10% to 50% and discarded the first two drive frequency intervals (instead of just the first) to determine their zero-frequency relative increase rate and maximum drive frequency. We used a linear fit for the data where r2 resistance allele frequency was between 0 and 0.5 with the same procedure.

### 2.6 Evaluation of chasing

We divided the simulation outcomes into three initial categories—population suppression without chasing, long-term persistence, and drive loss without chasing. After chasing starts, the population may finally be suppressed, or all drive alleles could be lost. However, it is also possible that drive, wild-type, and resistance alleles coexist for an extended period of time (until the end of the simulation). We hypothesized that in each chasing generation there is the same probability to eliminate the population or lose all drive alleles under a certain parameter set, which means that the time of chasing before suppression or drive loss would comply with a negative binomial distribution.

Because chasing is characterized by strong regional clustering of individuals, we used the average nearest neighbor ratio (Chester 1987) to quantify the degree of clustering during chasing. For each time step *j*, its average nearest neighbor distance 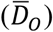 is defined as the average distances between each individual and its nearest neighbor, expressed as 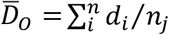, where *d*_*i*_ represents the distances between individual *i* and its nearest neighbor. The expected average nearest neighbor distance of time step *j* is the average nearest neighbor distance when individuals are randomly distributed and is calculated as 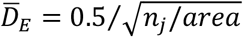, where *n*_*j*_ is the number of individuals. The average nearest neighbor ratio of time step *j* is given as 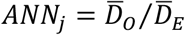 (**Figure 1B**). According to this definition, the average nearest neighbor ratio is smaller if individuals are more clustered.

Due to individuals tending to spread out slightly nonrandomly to avoid competition, a natural population is shifted toward a uniform distribution with equal spacing between individuals before any gene drive activity, so the average nearest neighbor ratio is a little higher than 1. The minimum value for *ANN* is 0, indicating that all individuals are at a single position. We found that the distribution of individuals and the value of average nearest neighbor ratio fluctuate nonrandomly around the average in chasing dynamics, with only small changes between time steps (**Figure 1C**).

If the average nearest neighbor ratio of a chasing generation is higher, it has more equilibrium character, which means that the local population density is approximately uniform across overall space. To provide a single value to characterize chasing across many generations for a specific simulation, we used a *weighted ANN* to moderate the influence of extreme values caused by time steps with few individuals. We defined this as *weighted ANN* = ∑_*time step*_ *ANN*_*j*_ * *n*_*j*_, including all time steps after chasing started. Note that this weighted average nearest neighbor ratio is more precise when the simulation is ended with long-term chasing, because short-term chasing dynamics include fewer time steps, and most of them contain fewer individuals, which increases the error of the *weighted ANN* of these simulations.

Chasing dynamics can also be characterized by the number of local chasing clusters. There is usually more than one group of wild-type individuals involved in chasing in a simulation at any time point (**Figure 1A**). The spatial distribution of these detected chasing cells (see above) contains information on the number of local chasing clusters. We used a clustering method based on a Density Based Spatial Clustering of Applications with Noise (DBSCAN) algorithm to cluster detected chasing cells and estimate the number of local chasing clusters. DBSCAN is effective in discovering clusters of arbitrary shape and does not need to predetermine the number of clusters (Ester *et al*. 1996).

DBSCAN determines clusters by finding “core points” that have at least a minimum number (*MinPts*) of neighbors within a given radius (*Eps*). The *Eps*-neighborhood of a point *p* is {*q* ∈ *R*|*dist*(*p, q*) ≤ *Eps*}, where *dist*(*p, q*) is the Euclidean distance between points *p* and *q*. For a core point *q*, if *p* is within the Eps-neighborhood of *q, p* is density-reachable from *q*, or if there is a chain of points *p*_1_, …, *p*_*n*_, *p*_1_ = *p, p*_*n*_ = *q*, in which *p*_*i*+1_ is density-reachable from *p*_*i*_ for every two adjacent points, then *p* is also density-reachable from *q*. With these definitions, the algorithm starts with an arbitrary cell *p* and labels it as a “core point” or “noise point.” It then retrieves all points that are density-reachable from core point *p* to get cluster C and labels them as a “cluster point”. This process is repeated until all of the detected chasing cells have been processed. Finally, all remaining “noise points” are ignored. In this study, we set the *Eps* as √5 and *MinPts* as 4, which means a local chasing cluster must have at least 4 chasing cells. However, because the local chasing detection method only detects expanding cells, this method loses efficiency when the population spreads to the boundary of the space. To address this, we set the *Eps* to 4 if two local chasing clusters have points on the same boundary.

### 2.7 Data generation

All simulations were implemented in the SLiM forward-in-time population genetic simulation framework, version 4.0.1 (Haller & Messer 2023). Data processing and analytics were performed in Python. All code and data are available on GitHub (https://github.com/xinga1516/chasing).

## 3 Results

### 3.1 Weighted ANN

We first sought to better quantitatively characterize chasing after the release of a homing suppression gene drive. The average population size or number of fertile females is an important measure, but it does not itself have information about the distribution of individuals. We therefore utilized the average nearest neighbor (ANN) distance to quantify this distribution. It has been shown that both the drive conversion and female heterozygote fitness, which is often reduced by somatic expression of Cas9, have a significant influence on the exact level of suppression from the female fertility drive system (North *et al*. 2020; Champer *et al*. 2021a; Champer *et al*. 2022; Liu & Champer 2022). Here, we used these two parameters to test the performance of our chasing assessment method (**Figure 2**). Though population elimination without chasing could occur for highly efficient drive (**Figure 2A**) it would often be accompanied by chasing dynamics (**Figure 2B**). Over a large region of parameter space, the drive would persist, but fail to eliminate the population (**Figure 2C**). We recorded the suppression rate per generation during chasing (**Figure 2D**), as well as the average population size during long-term persistence of both drive and wildtype alleles(**Figure 2E**).

**Figure 2.**
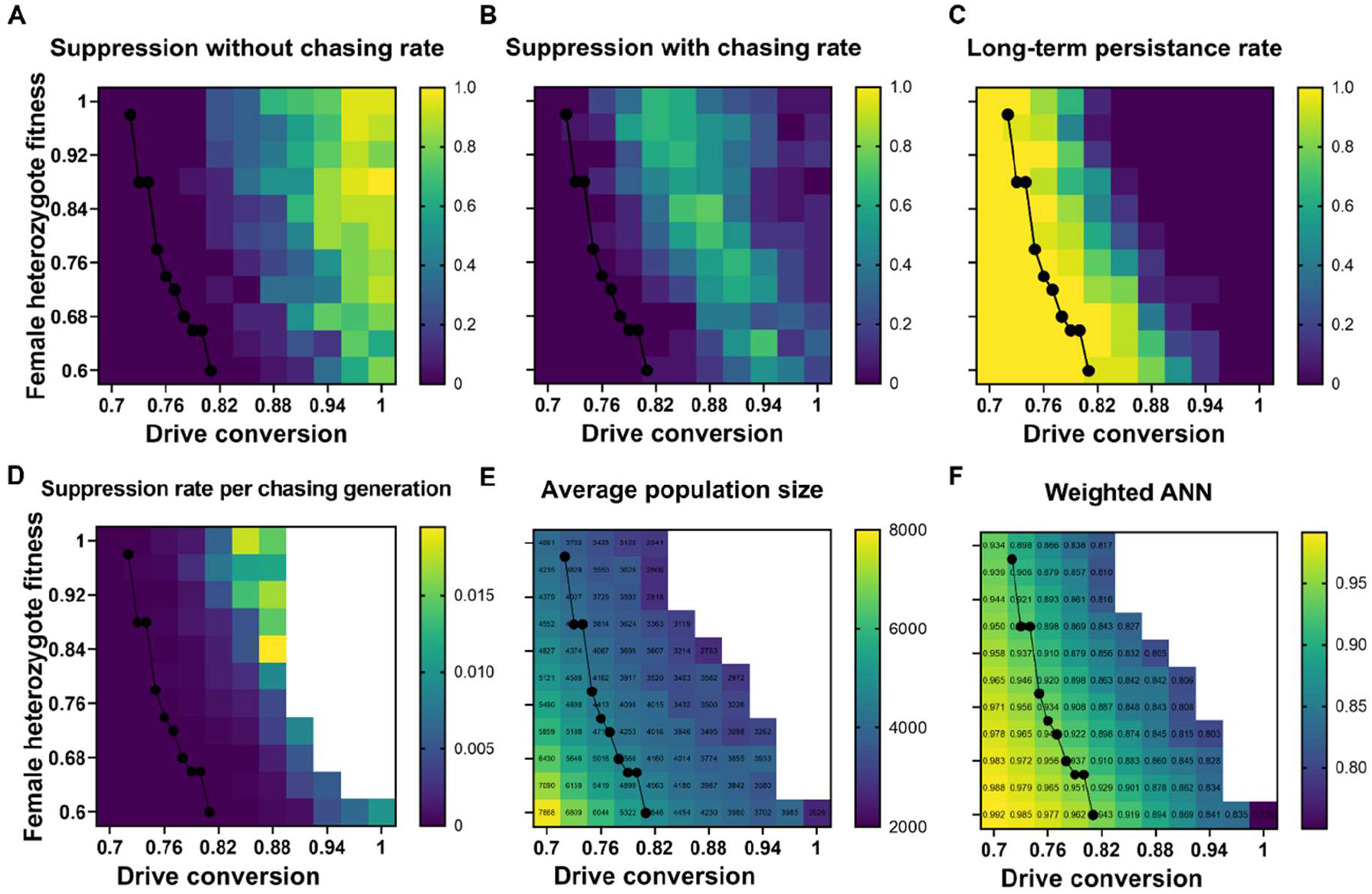
Characterizing of chasing outcomes. Simulations use the discrete-generation fecundity model with 20 simulations for each combination of varying drive conversion and female heterozygote fitness. Other parameters are at their default values. Heatmaps show rates of (**A**) population elimination without chasing, (**B**) population elimination after chasing, and (**C**) long-term persistence of both drive and wild-type alleles. Also displayed are (**D**) probability each generation during chasing that the population is eliminated and (**E**) the average long-term population size, and (**F**) the average *weighted ANN* during long-term persistence. The black lines in the heatmaps show the threshold values that produce a sufficient genetic load to eliminate a panmictic population.

The *weighted ANN* presented the differentiation of all these chasing dynamics under different parameter combinations (**Figure 2E**). The *weighted ANN* is lower when the drive is more powerful, with a higher drive conversion rate and female heterozygote fitness, which corresponds to a higher suppression with chasing rate and a higher suppression rate per chasing generation. This is consistent with the fact that a powerful drive can make the population clustered enough to completely eliminate it. When the drive conversion is 0.7 and the female heterozygote fitness is 0.6, representing a less powerful drive system, the value of the *weighted ANN* is close to 1, which means that the population is in an approximately random distribution. This is regarded as an equilibrium scenario in former studies (Champer *et al*. 2021a).

If the parameters are insufficient to eliminate the population in a panmictic model (the black lines in **Figure 2**), then population suppression will always fail in the spatial model. This is consistent with former models (Eckhoff *et al*. 2017; North *et al*. 2020; Champer *et al*. 2021b; Birand *et al*. 2022). In addition, population size during chasing is correlated with the *weighted ANN*, and the points of the genetic load threshold line (showing where the drive has enough suppressive power to eliminate a similar panmictic population) almost fall on a narrow range of *weighted ANN*. Because all these threshold points have the same genetic load, it is potentially reasonable to extrapolate that the *weighted ANN* represents the drive power under certain ecological parameters, and it can give an effective assessment of the drive performance under different scenarios to some extent. The continuous nature of the *weighted ANN* also shows that there is not a clear boundary between chasing and equilibrium outcomes. When the genetic load is greater than the threshold, the outcome can certainly be considered to be chasing if rapid population elimination fails. Below this threshold, the outcome in panmictic populations is equilibrium, but because spatial models have greater local fluctuations, local suppression will still often occur, followed by recolonization. Slightly below the threshold line, these dynamics are essentially the same as slightly above the line, being recognizable as “chasing.” Eventually though, as the *weighted ANN* approaches 1, the outcome smoothly transitions into something more easily recognizable as equilibrium, with little to no local stochastic elimination.

### 3.2 Chance per generation of population elimination

If chasing represents a dynamic equilibrium state, then after this equilibrium is reached, we hypothesized that in each chasing generation, there is the same probability to eliminate the population (or lose all drive alleles) over a sufficiently large time interval, which means that the time of chasing before suppression or drive loss would comply with a negative binomial distribution. To test this, we ran 100 simulations using the discrete-generation fecundity model and collected outcomes and the time that chasing was initiated. We conducted a goodness of fit test to check if the time that chasing ends complies with a negative binomial distribution. The *p*-value of the Pearson Chi-square test was 0.778, meaning that the negative binomial distribution can moderately well represent the distribution of the duration of chasing. Thus, if we can run sufficient simulations with outcomes ending in successful population elimination or drive loss, we can estimate their probability per chasing generation by maximum likelihood estimation. Though the suppression rate of a certain generation is dependent on the population size and spatial distribution, the chance of population elimination per time step is still a good way to assess suppression outcomes during chasing.

### 3.3 Effect of arena size

In most models of chasing in continuous space, the arena size represents only a small fraction of the landscape in many relevant real-world scenarios. This is by necessity due to computational resource limitations. However, large arenas can potentially drastically change the outcomes. Here, we analyzed chasing dynamics in spaces with different arena sizes using the same population density. **Figure 3** shows that the arena size has a great influence on chasing dynamics with the same drive system. As the arena size increased, the probability of chasing also increased (**Figure 3A**). After chasing started, suppression and drive loss rates per generation exponentially declined with a linear increase in arena size (**Figure 3B**). To explain this result, we computed the *weighted ANN* and clustered local chasing cells in each time step. We found that the *weighted ANN* gradually leveled off at 0.67 with increasing arena size (**Figure 3C**), and the average number of local chasing clusters and the average population size are both almost directly proportional to arena size, resulting in the average population size of each chasing cluster gradually leveling off at 4,740 (**Figure 3D**).

**Figure 3.**
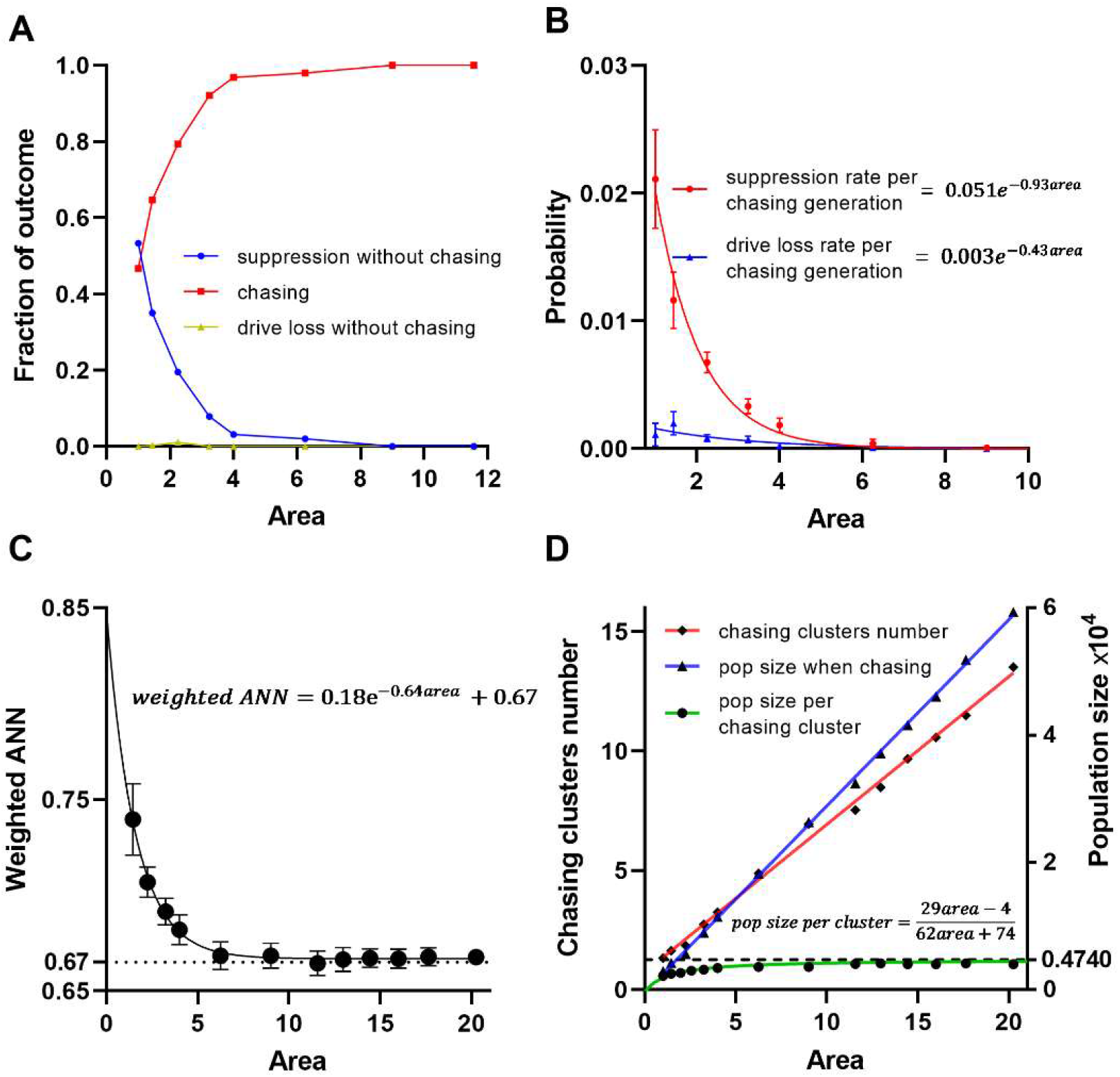
Effects of the arena size on suppression outcomes and chasing dynamics. Using the discrete-generation fecundity model, we set the drive conversion to 0.94 and the population density to 10,000, with other parameters at default values. (**A**) The fraction of three possible drive outcomes with at least 40 simulations for different arena area. (**B**) The rate of population elimination or drive loss per generation once chasing starts. The error bars are 95% confidence intervals of the estimated success probability from the Bernoulli distribution. Data is fit with the exponential function. (**C**) The weighted ANN as a function of area with 40 simulations that avoided population elimination or drive loss per parameter set. Error bars are 95% confidence intervals, and the data is fit with an exponential function. (**D**) The population size and number and size of chasing clusters. The population size per chasing cluster is based on the linear regression lines for number of clusters and total population size.

Based on the number of local chasing clusters (*n*) and the population size per local chasing cluster (*s*), the probability of suppressing a local chasing cluster should be a function of *s* and the aggregation degree of individuals — *weighted ANN*. Lower *weighted ANN* should result in more clustered local chasing with a higher probability of population elimination. Smaller chasing clusters should be easier to eliminate. Thus, the elimination rate per chasing generation can be expressed as [*f*(*s, ANN*)]^*n*^. Because *n* increases linearly with the size of the arena for sufficiently large arena size, the suppression rate per chasing generation should exponentially decline with increasing arena size, which is consistent with our observations. The same conclusion applies to an assessment of drive loss rates.

When the arena is smaller, *weighted ANN* is higher, which means that chasing dynamics tend to be less extreme in terms of the uneven distribution of individuals. This may be due to edge effects on moving chasing clusters, which will be far more impactful in a small arenas. When the arena is large enough for the population size per chasing cluster and *weighted ANN* to level off, the number of chasing clusters would proportionally grow with the arena size, but the global distribution would no longer change significantly.

Hence, as arena size increases, *f*(*s, ANN*) levels off, so the suppression rate per chasing generation will rapidly decrease to 0. In other words, if the area is large enough, once chasing dynamics starts across the entire arena, there is little chance for it to stop. If any local clusters are eliminated, the system will quickly recover because many other clusters will still be present. The only way to successfully eliminate the population under these circumstances in the model would be to prevent chasing from starting in the first place.

### 3.4 Resistance allele formation

Resistance alleles have been generally regarded as an important factor that can lead to the failure of suppression drive, especially when the drive has a high fitness cost (Price *et al*. 2020; Hammond *et al*. 2021). While functional r1 resistance alleles will cause rapid drive failure, nonfunctional r2 resistance alleles can also slow the drive (though they will have only a modest effect on genetic load if other drive performance parameters are good). Yet these have not been assessed for chasing situations. There are two ways to generate r2 resistance alleles in CRISPR gene drive systems. One of them involves end-joining repair after cleavage in the germline as an alternative to homology-directed repair. We varied the germline resistance rate and found that the probability of chasing increases with the germline resistance rate (**Figure 4A**). Further, the suppression rate of each generation during chasing exponentially declines with increasing germline resistance rate (**Figure 4B**).

**Figure 4.**
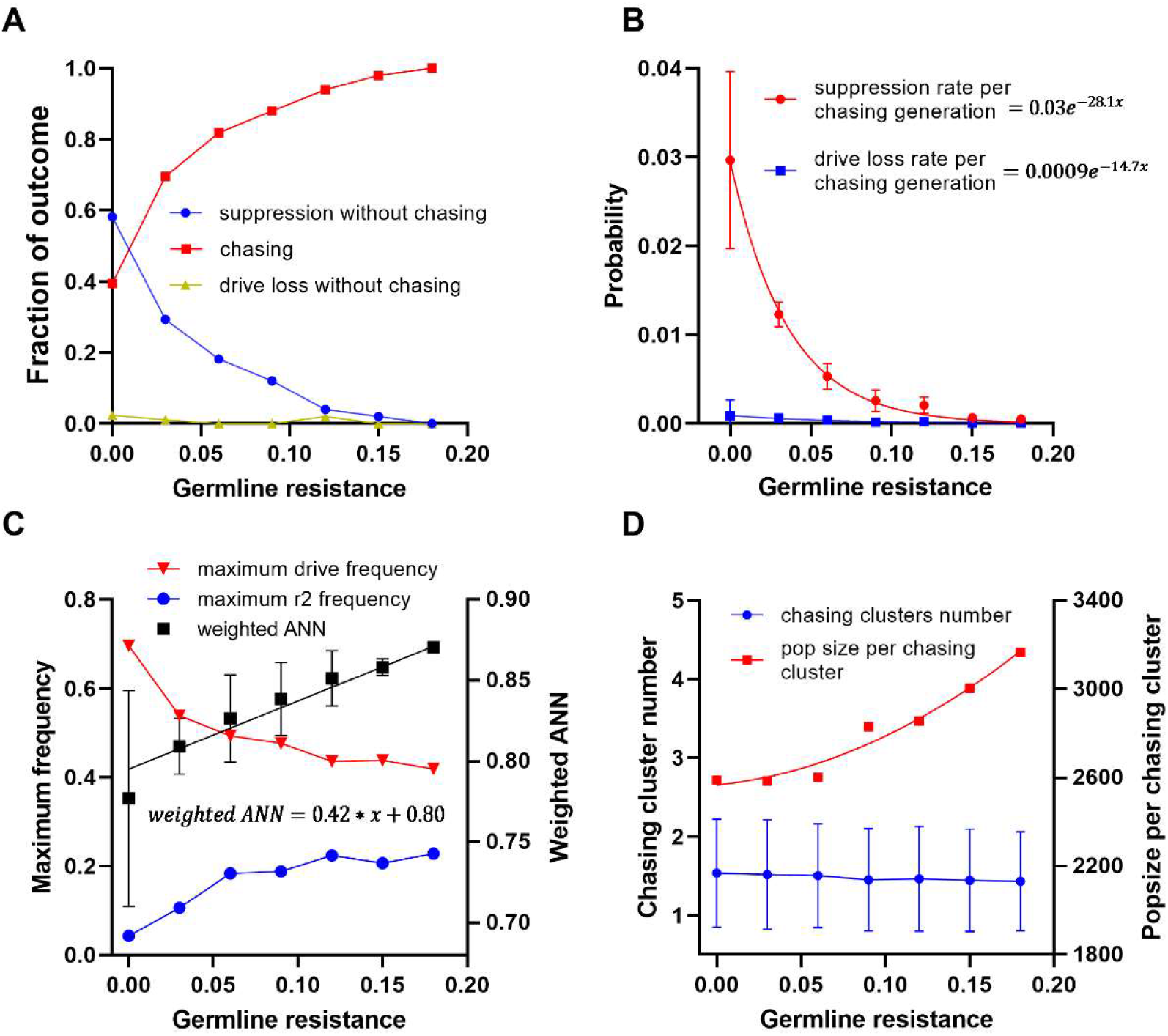
Effects of germline resistance rate on suppression outcomes and chasing dynamics. Using the discrete-generation fecundity model, we set the drive conversion to 0.82, and other parameters were at default values. (**A**) The fraction of three possible drive outcomes with at least 50 simulations for each different germline resistance rates. (**B**) The rate of population elimination or drive loss per generation once chasing starts. The error bars are the 95% confidence intervals of estimated suppression rate per chasing generation of a Bernoulli distribution. Data is fit with the exponential function. (**C**) The weighted ANN as a function of germline resistance rate (error bars are 95% confidence intervals), as well as the maximum drive and drive frequency at equilibrium. (**D**) The population size per chasing cluster and number of chasing clusters. Error bars show standard deviation.

To learn about the mechanism of how nonfunctional resistance affects chasing, we examined how they hinder the spread of drive. We determined the drive and r2 allele relative increase rate (**Figure S3**) and their maximum frequency (**Figure 4C**). The zero-frequency relative increase rate of drive alleles represents the drive’s relative rate of growth when drive frequency is very low (for example, an ideal homing drive will have a relative growth rate of 2, because the whole population will be drive/wild-type heterozygotes that double the drive frequency in the next generation). This measure has little change with increasing germline resistance rate, while the relative increase rate of r2 alleles grows. With the same zero-frequency relative increase rate of drive alleles, the lower maximum drive frequency means a more dramatic decline of the relative increase rate, manifesting the lack of power to increase when frequency is high. This alone has little effect on genetic load because the increased number of nonfunctional resistance alleles at equilibrium allow for nearly the same total number of recessive sterile alleles (**Figure S4**). However, the ability of the drive to migrate from high frequency to low frequency is hindered, which may be important when considering spatial spread during chasing. Because the number of local chasing clusters shows little difference (**Figure 4D**), this appears to manifest as higher *weighted ANN* (**Figure 4C**) and higher population size per chasing cluster (**Figure 4D**), reducing the probability for a local chasing cluster to be eliminated.

Another way for resistance alleles to be formed is when wild-type alleles in early embryos are cut by maternally deposited Cas9 and gRNA. Unlike germline resistance allele formation, embryo resistance can only be generated by females, but on the other hand its rate can vary widely without being constrained by the drive conversion rate (the drive conversion and germline resistance rates can together add up to no more than one). We performed a similar analysis of the effect of embryo resistance alleles on chasing and found that over the full parameter range, it could have an even larger effect reducing successful suppression and increasing chasing (**Figure 5A-B**). Because the initial rate of decrease in the suppression rate is slower, a Gaussian function curve better fits the suppression and drive loss rate per chasing generation, which still rapidly declined with increasing embryo resistance. The *weighted ANN* and population size per chasing cluster similarly increases to a greater degree (**Figure 5C-D**), which can explain why the chance per generation of population elimination dramatically decreased.

**Figure 5.**
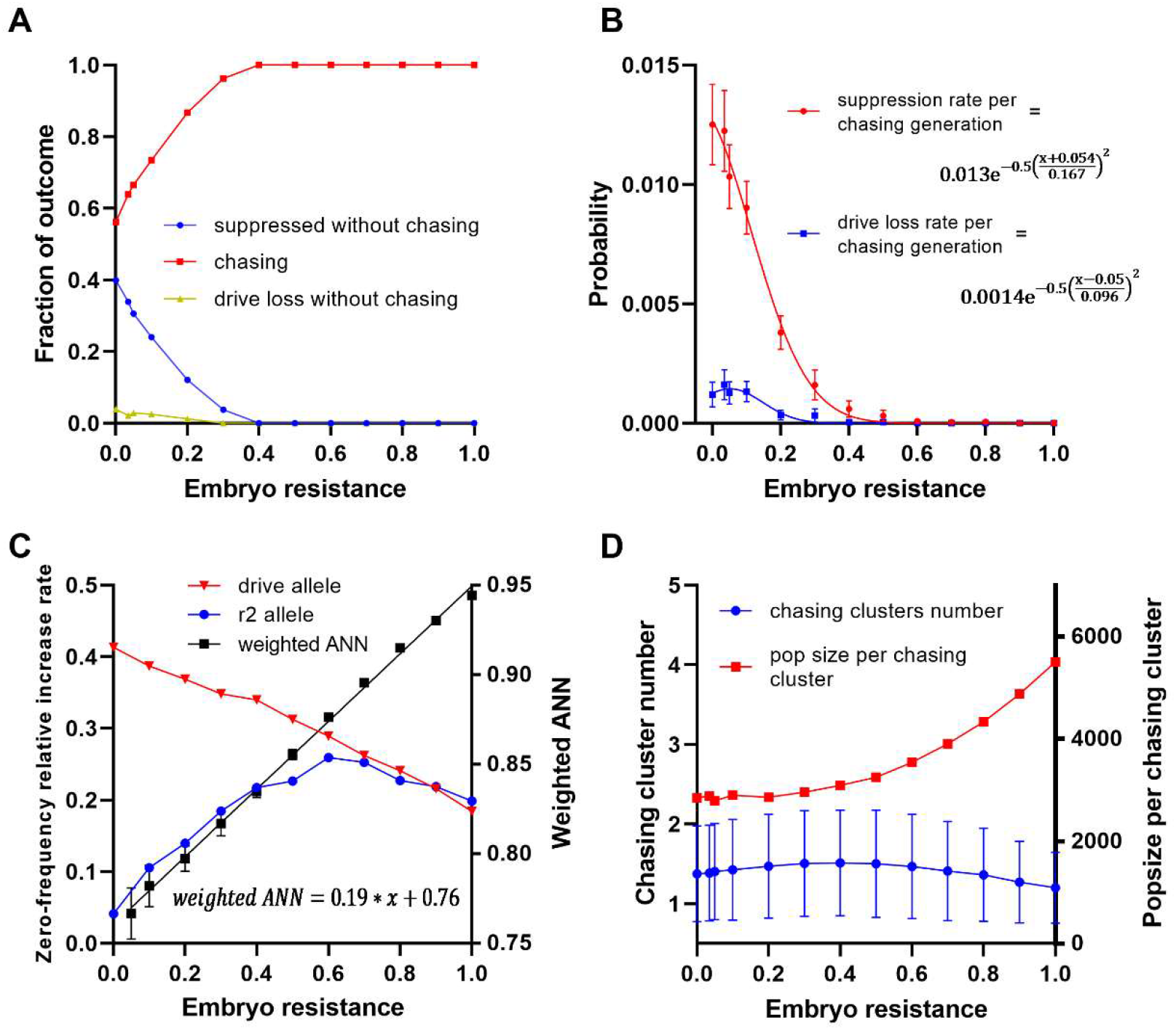
Effects of embryo resistance rate on suppression outcomes and chasing dynamics. Using the discrete-generation fecundity model, we set the drive conversion to 0.88, and other parameters were at default values. (**A**) The fraction of three possible drive outcomes with at least 60 simulations for each different embryo resistance rates. (**B**) The rate of population elimination or drive loss per generation once chasing starts. The error bars are the 95% confidence intervals from a Bernoulli distribution. Data is fit with the exponential function. (**C**) The *weighted ANN* as a function of embryo resistance rate (error bars are 95% confidence intervals), as well as the maximum drive and drive frequency at equilibrium in a panmictic population. (**D**) The population size per chasing cluster and number of chasing clusters. Error bars show standard deviation.

One additional explanation for this greater effect could be that unlike germline resistance, increasing embryo resistance rate can slow down the initial drive spread speed (**Figure 5C, Figure S4**). This can be understood when considering that at low drive frequencies mating of drive heterozygote females will usually be to wild-type males. Germline alleles formed during this process will not affect the drive, but embryo resistance alleles that form in progeny with a drive allele will result in sterile females, removing the drive allele from the population. At drive higher frequencies, any resistance allele (such as one formed in the germline) is more likely to eventually meet a drive allele and be removed from the population. Thus, embryo r2 resistance alleles can directly slow the increase speed of drive alleles, which will slow the drive wave advance speed. This will leave more time for wild-type individuals to escape and recover, perpetuating chasing cycles and explaining the larger population size during chasing with high embryo resistance (**Figure 5D**). When embryo resistance is over approximately 0.5, the relative drive increase rate at low frequency continues to decline linearly, but the relative r2 increase rate also starts to decline (**Figure 5C**), reducing the suppressive effect on the population. Because the r2 resistance alleles are nonfunctional, they tend to be eliminated in female progeny together with drive alleles, eventually reducing the rate that both increase and accounting for the reduced rate of drive increase.

### 3.5 Effect of life history traits on chasing dynamics

Previous studies indicated that chasing was a substantially larger problem for gene drives in a mosquito-specific model with overlapping generations than a simpler discrete-generation model (Champer *et al*. 2022). The mosquito model allowed a small fraction of females to remate in each time step, but in the discrete-generation model, a female would only have one mate when producing all offspring. These models also differed in the way in which population size was regulated. In the mosquito model, there was density-dependent competition between juveniles, affecting larval viability, while in the discrete-generation model adult density-dependent competition affected female fecundity. To investigate how such life history details of a species can influence chasing dynamics, we implemented both kinds of reproduction strategies (viability and fecundity) in both a discrete-generation model and an overlapping-generations model.

The drive performance in species with overlapping generations and discrete generations was substantially different when females were allowed to remate in each time step. We found that the overlapping-generations model produced fewer chasing outcomes and slightly fewer drive loss without chasing outcomes (**Figure 6A**). During chasing, the suppression rate per generation was also higher for overlapping generations (**Figure 6B**). This is because when chasing starts, the overlapping generations model has smaller and more clustered local chasing clusters than the discrete-generation model (**Figure 6C-D**).

**Figure 6.**
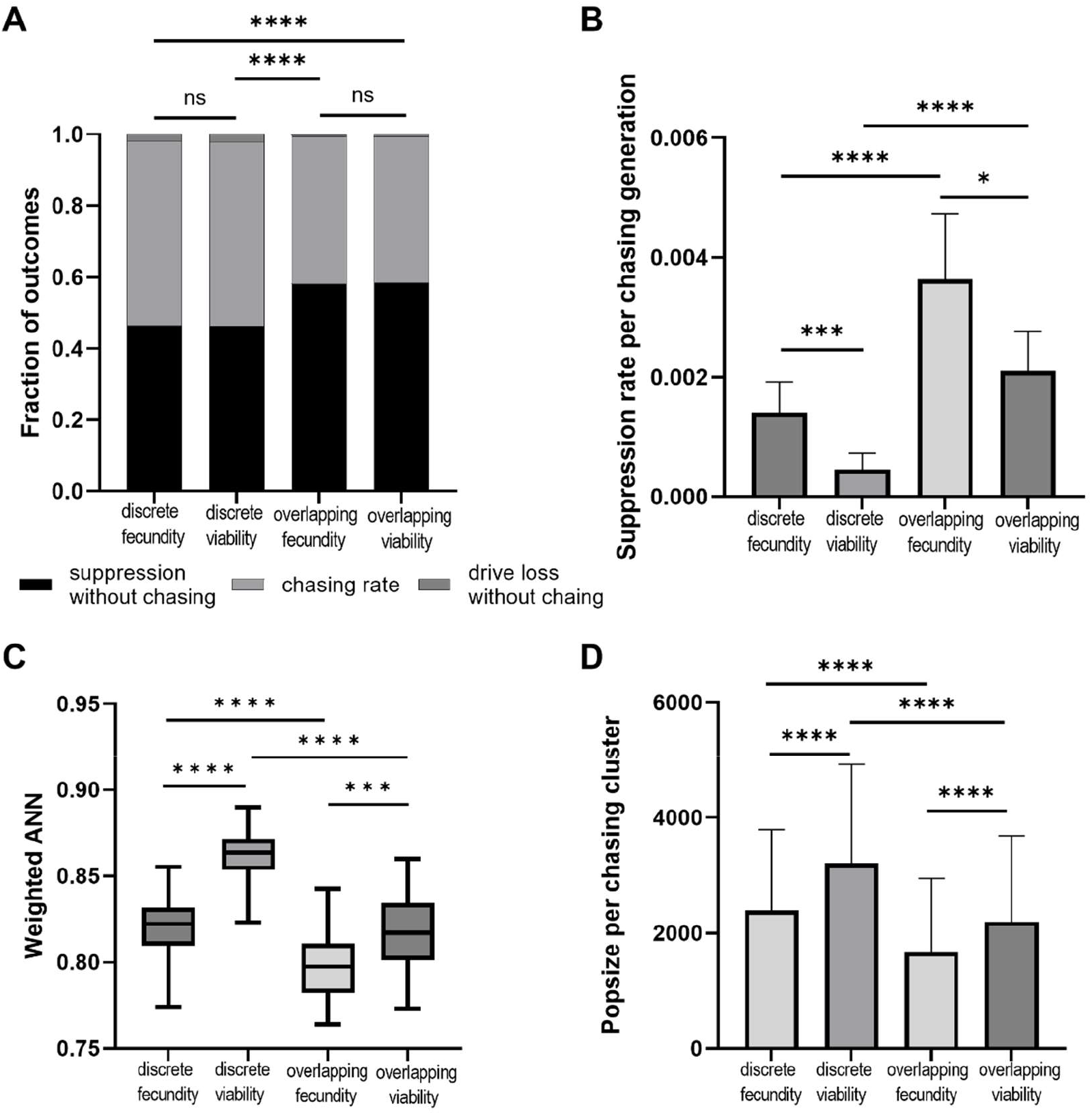
The differences of chasing dynamics between model types. We compared discrete-generation and overlapping-generations models, and also fecundity and viability models. (**A**) The drive conversion is 0.92, and other parameters are default value. The fraction of outcomes with 100 simulations is shown. (**B-D**) The drive conversion is 0.86, and the female heterozygote fitness is 0.8. Other parameters are at default values. (**B**) The suppression rate per generation during chasing. The error bars are 95% confidence intervals from a Bernoulli distribution. (**C**) The *weighted ANN* from each model (error bars are 95% confidence intervals). (**D**) The median population size per chasing cluster (error bars show interquartile range). Statistical comparisons are the Chi-square test for **A-B** and unpaired t-test for **C-D**. * *p* < 0.05, ** *p* < 0.01, *** *p* < 0.001, **** *p* < 0.0001.

In terms of outcomes without chasing, there was no difference between the viability and fecundity models (**Figure 6A**). However, viability-based models consistently had lower suppression rates per chasing generation, as well as less clustering and larger population sizes per chasing cluster (**Figure 6B-D**). Due to greater stochasticity, drive loss was also higher in fecundity models (**Figure S5**). Because the influence of overlapping generations and viability competition was in opposite directions, the difference of the mosquito model remained unexplained by these factors alone.

To further investigate different life histories, we made an overlapping-generations model in which females that had successfully mated would store sperm of the first mate and reproduce at each time step, but not mate again. This was closer to the mosquito model, in which females only had a 5% chance to remate in each weekly time step, in line with field data (Champer *et al*. 2022). We found that the suppression rate per chasing generation reached the same level as in the discrete-generation model when we prevented remating (**Figure 7A**). In addition, the *weighted ANN* was even lower, and the population size during chasing grew to a higher level than in the discrete-generation model (**Figure 7B-C**). Therefore, remating is an important reason why the suppression rate per chasing generation rose in the overlapping-generations model. Effects on drive loss were minor (**Figure S6**).

**Figure 7.**
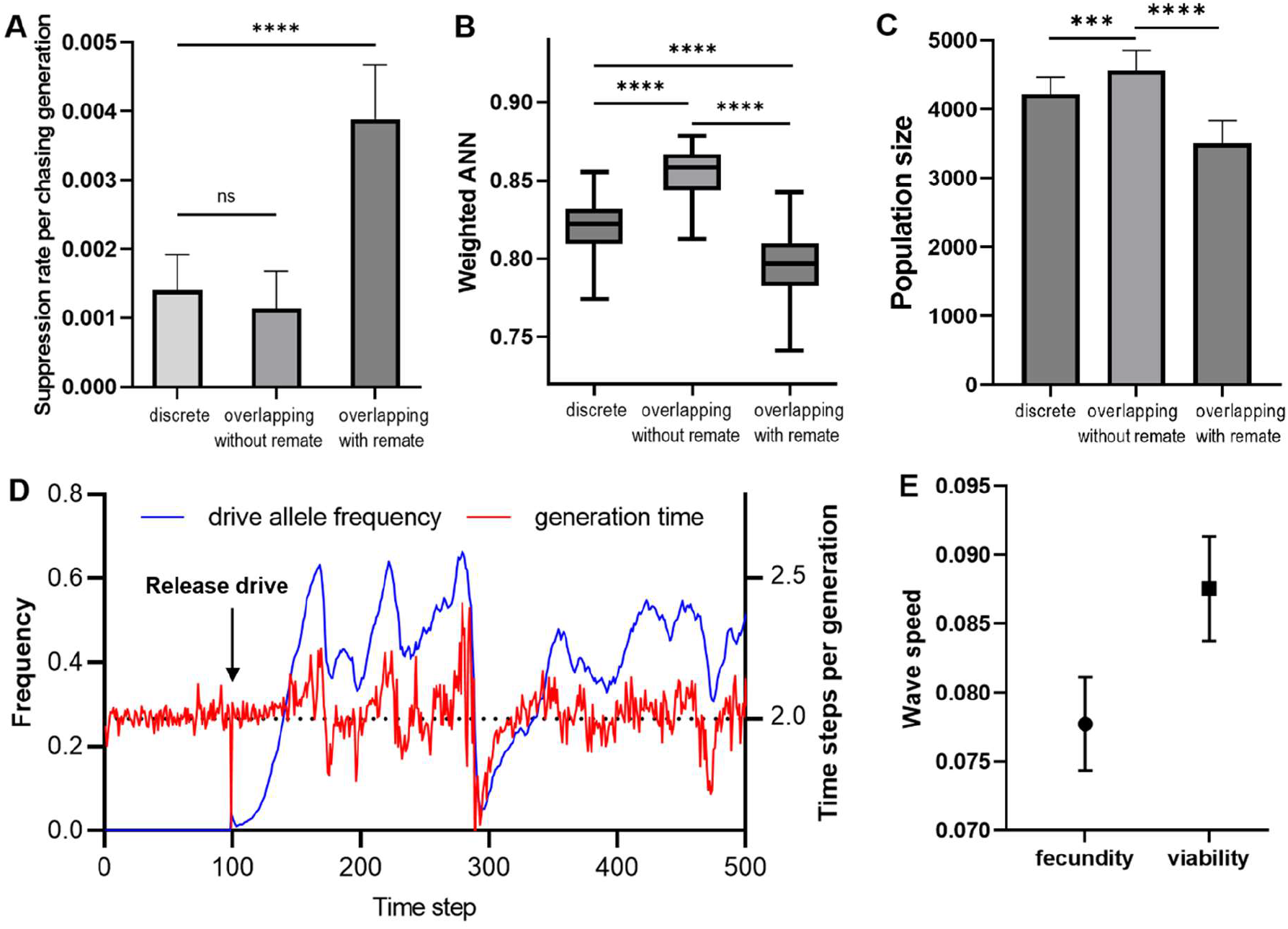
The influence of remating on chasing dynamics. In the fecundity model, the drive conversion is 0.86, and the female heterozygote fitness is 0.8. Other parameters are at their default values. (**A**) The suppression rate per generation during chasing (error bars are the 95% confidence intervals from a Bernoulli distribution). (**B**) The *weighted ANN* during chasing (error bars are 95% confidence intervals). (**C**) The median total population size during chasing (error bars show the interquartile range). (**D**) The change of drive frequency and time steps per generation in one simulation of the overlapping-generations model without remating. The generation time in theory is two time steps. (**E**) The drive wave speeds in the fecundity model and viability model. The mean and standard deviation are shown from 20 replicates. Statistical comparisons use the Chi-square test for suppression rate per chasing generation, and unpaired t-test for *weighted ANN* and *population size per chasing cluster*. ns: not significant, * *p* < 0.05, ** *p* < 0.01, *** *p* < 0.001, **** *p* < 0.0001.

However, remating is not the only factor that caused the differences between the discrete-generation model and overlapping-generations model. The generation time (calculating the average reproduction age of all fertile females weighted by their offspring numbers) in the overlapping generations model without remating is usually two time steps with low variation (**Figure 7D**). During chasing, though, it fluctuates, largely based on drive frequency. The generation time grows during the suppression process but shortens during the recovering process for wild-type individuals (**Figure 7D**), thus promoting chasing and reducing the chance of suppression. Younger females produce more offspring when competition is low, such as at the edges of a chasing cluster, and lower generation time allows faster recovery of escaped wild-type individuals. These also influence chasing dynamics and can partly explain why the average population size during chasing is higher and the clustered degree is lower in the overlapping-generations model without remating than in the discrete-generation model (**Figure 7B**-**C**). This property causes the wild-type wave advance speed to become faster than in the discrete-generation model, which may further explain why chasing is more clustered in the overlapping-generations model with higher cluster sizes. What’s more, we found the wild-type wave advance speed into empty space of the viability model is higher than the fecundity model (**Figure 7E**). This is because in the viability model, the survival rate of offspring at the edges of the population clusters is higher than the offspring in the middle, which helps the spread of wild-type individuals into empty space. In contrast, the positions of offspring are set when they are born in fecundity model, where even females closer to the edge have more competition than widely dispersing offspring.

All these factors can explain the difference in chasing dynamics between the different models. Overall, overlapping generations would hypothetically allow for less chasing in the mosquito model (Champer *et al*. 2022), but this is counteracted by lack of remating. Then, the viability verses fecundity density-dependent competition type difference can explain why chasing is a more common outcome in the mosquito model.

### 3.6 Density growth curve shape

With this study on different life history models and a previous study on competing species (Liu *et al*. 2023) showing the importance of ecology on chasing dynamics, we were interested in investigating the influence of different density-dependent growth curves (**Figure 8A**).

**Figure 8.**
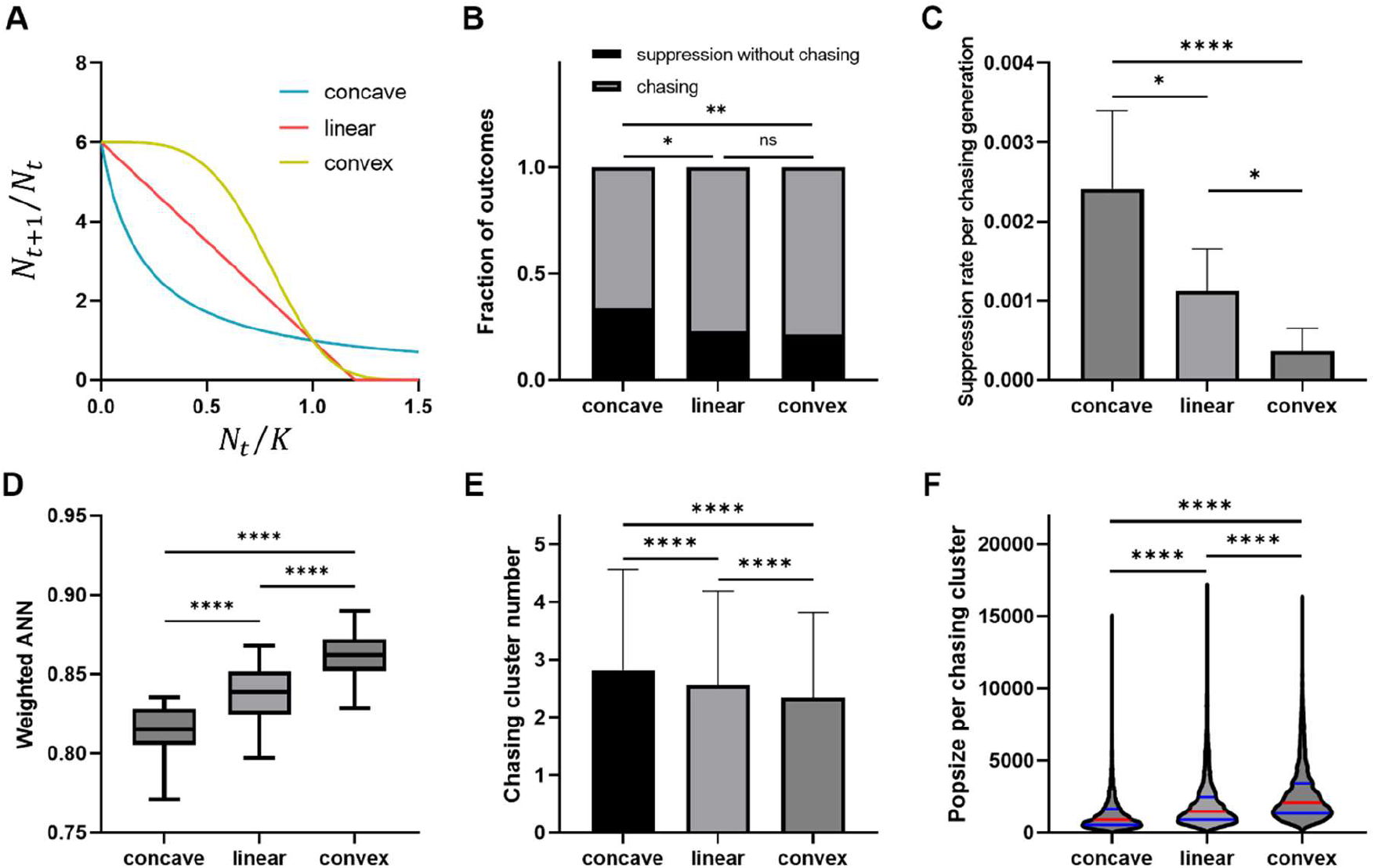
Effects of the density growth curve shape on suppression outcomes and chasing dynamics. (**A**) The three different density growth curves used in this study are displayed with a low-density growth rate of 6. In the overlapping-generations fecundity spatial model, (**B**) the fraction of chasing and suppression without chasing outcomes are shown for different density growth curves when the female heterozygote fitness is 0.8, and other parameters are at their default value. There are 200 simulations for each parameter set. (**C-F**) In the overlapping-generations fecundity spatial model, drive conversion is 0.84, female heterozygote fitness is 0.8, and other parameters are at their default value. (**C**) The suppression rate per generation is shown with different density curve shapes (error bars are the 95% confidence intervals of the estimated success probability with a Bernoulli distribution). (**D**) The *weighted ANN* from each density curve shape (error bars are 95% confidence intervals). (**E**) The chasing cluster number from each density curve shape (error bars are standard deviations). (**F**) A violin plot of population size of each chasing cluster. The red lines represent the median value of the data, and the blue lines are quartiles. Statistical comparisons are the Chi-square test for **B-C** and unpaired t-test for **D-F**. ns: not significant, * *p* < 0.05, ** *p* < 0.01, *** *p* < 0.001, **** *p* < 0.0001.

Convex, and to a lesser extent linear, density curves will tend to produce more robust populations. However, this would generally only delay and not prevent population elimination in panmictic populations, which is determined by whether the genetic load of the drive can overcome the low-density growth rate (though increased stochasticity could also slightly ease the requirement for elimination in concave curves for smaller population sizes). The magnitude of the density curve shape effect on chasing is unclear. In general, concave density growth curves resulted in less chasing dynamics than linear and convex curves (**Figure 8B**). However, the suppression rate per chasing generation (**Figure 8C**) was more strongly affected, with concave curves producing significantly higher population elimination rates than linear curves, and convex curves producing significantly lower elimination rates.

The *weighted ANN* (**Figure 8D**) was similarly affected, with more population clustering with a concave density curve, and fewer in convex. The concave density-dependent growth curve results in more chasing clusters with smaller population size (**Figure 8E**,**F**). In the convex and to a lesser extent, linear curves, when the population size starts to decline from equilibrium, the growth rate increases more quickly than concave, allowing faster recovery. In a panmictic setting, the drive will maintain pressure on the population, preventing recovery. A different growth curve can change the time to population elimination, but it won’t change the ultimate outcome, except indirectly via stochastic effects. However, chasing in a spatial model will have wild-type individuals that can take advantage of the faster recovery from linear and convex curves. When wild-type individuals escape from drive individuals and reach the empty space, populations with convex density-dependent growth curves will have higher growth before drive individuals reach them. This may explain why convex density-dependent growth curves had greater population size per chasing cluster and *weighted ANN*.

We also tested the effects of density-dependent growth curves on a larger range of parameters, including variation in the low-density growth rate with three different density-dependent growth curves using the overlapping-generations fecundity model. Drive conversion was also co-varied with low density growth rate in each case. Examining rates of long-term chasing, we found that concave density-dependent growth curves cause less chasing across the full parameter space (**Figure 9A**). The frequency of long-term chasing increases with the low-density growth rate, which is consistent with previous results (Champer *et al*. 2021a). Higher low-density growth rate allows more rapid wild-type propagation, which makes it easier to escape from the drive and recover. Species with linear or convex density-dependent growth curves were more prone to long term chasing, requiring more powerful suppression drives to successfully eliminate the population (**Figure 9B-C**). There was little change between linear and convex. Examining the duration of all chasing outcomes, linear and especially convex density-dependent growth curves had longer times when the low-density growth rate was sufficiently high (**Figure S7**).

**Figure 9.**
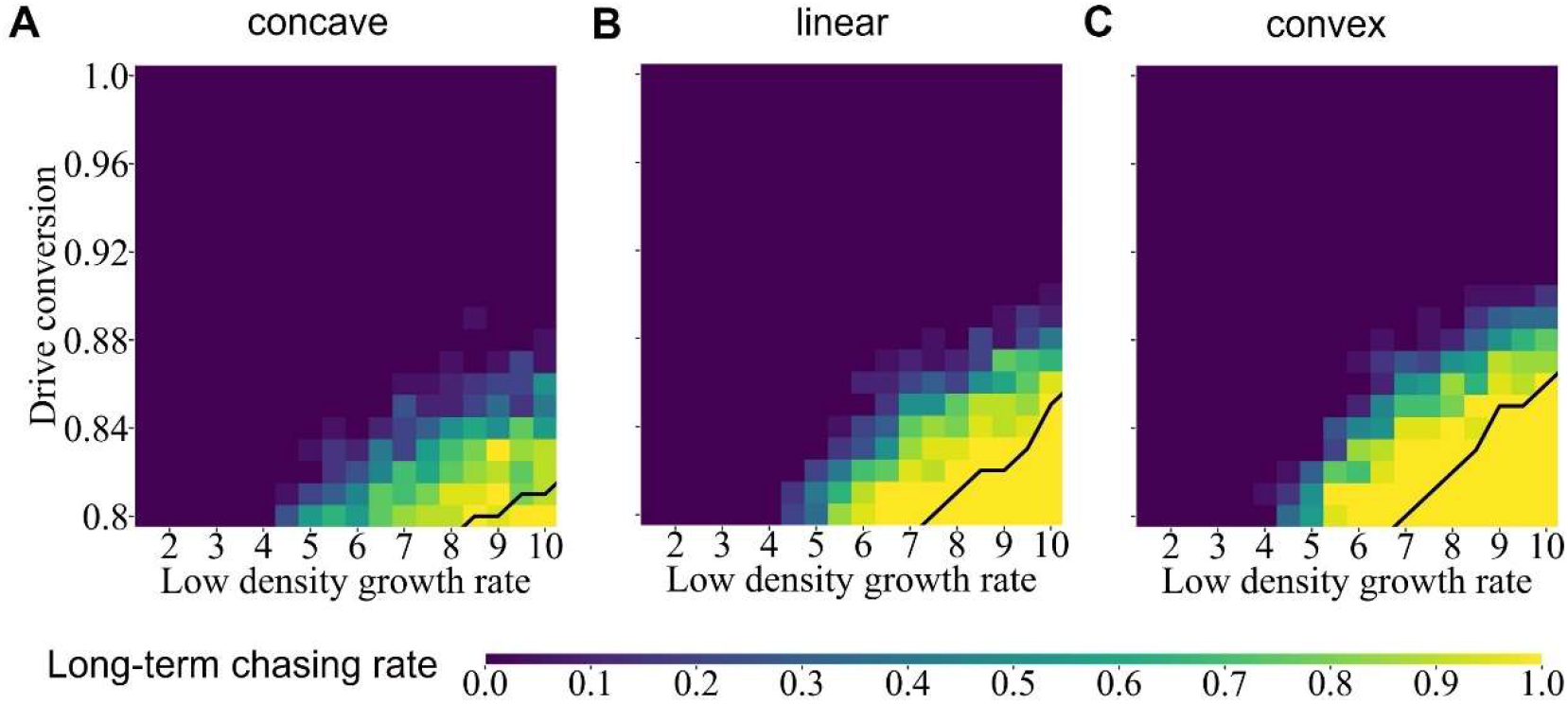
Long-term chasing rates with different drive performance and density growth curves. In the overlapping-generations fecundity model with a population density of 50,000 and other parameters with their default values, long-term chasing outcomes are show for varied drive conversion and low-density growth. Displayed are (**A**) concave, (**B**) linear, and (**C**) convex density growth curves. Each parameter set has 20 replicates. The black line shows the threshold above which the genetic load is sufficient to eliminate the population in a panmictic model.

## 4 Discussion

In this study, we examined the chasing phenomenon whereby suppression drive can fail to eliminate target populations with spatial structure. Because previous definitions of chasing were qualitative in nature or suitable for only a limited parameter set (Bull *et al*. 2019; Champer *et al*. 2021a) we developed new methods to classify and characterize chasing. We found that there is no clear line between chasing and static equilibrium dynamics in spatial models. We thus defined *weighted ANN* to quantify spatial distribution of individuals, which provides a clear measure to characterize chasing and the final expected performance of the drive, together with population sizes.

In panmictic populations, a suppression drive will be able to reliably eliminate the population if its genetic load is sufficiently high. Otherwise, the outcome will be an equilibrium state, where the average population is usually reduced from its initial level. However, in spatial models, chasing can prevent population elimination even when the genetic load would be sufficient in a panmictic population. Classifying chasing is somewhat difficult, though, because there does not seem to be a distinct point to differentiate chasing and equilibrium outcomes in spatial models. Chasing is characterized by a large variance of population density across time and space. Because the variance is a continuum, if we set a threshold, it must be based on an objective criterion. While models that fail when the genetic load is above its panmictic threshold can certainly be classified as chasing, we could not determine a way to distinguish chasing from equilibrium when the genetic load is below this value. Furthermore, it may not even be possible to distinguish these outcomes based on simulations if the expected panmictic outcome is not known. This reinforces the need to characterize chasing and equilibrium outcomes by quantitative measurements such as the *weighted ANN*.

Considering that wild-type recolonization is an independent event from each local area, our local chasing detection and clustering method based on a grid defined by average dispersal provides a direct and intuitive measure of chasing in a local area. The clustering method based on DBSCAN resolves noise and allows characterization of the number of chasing clusters. The size of the detection cell determines the resolution of local dynamics that can be detected, and we chose the average dispersal as a universal measure representing an important small-scale distance. Competition distance could also have been a reasonable choice.

Using these methods, this study also provides a guide to experiments. We showed that for large regions (arena size), if chasing is possible, it is very unlikely to end. Thus, drive performance must be sufficiently high to avoid chasing if population elimination is required. Ecological factors such as competing species could somewhat ease this requirement (Liu *et al*. 2023), but our study also showed that even nonfunctional resistance alleles can be highly problematic for chasing and should be minimized. Alternatively, if some remaining population can be tolerated, then the functional resistance allele formation rate must be substantially lower than what would be predicted in a panmictic population because chasing substantially increases the chances that they could form (Champer *et al*. 2021a; Liu & Champer 2022).

Due to the low probability of drive loss, our conclusions for this outcome are less robust than for population suppression. However, it likely that the drive loss rates per chasing generation in the fecundity and viability models have a significant difference (**Figure S5**). More clustered and smaller scale local chasing not only increases the chance of successful population elimination, but also increases the chance of drive loss. This is another example where the chance of drive loss cannot be avoided if successful target elimination is also a possibility, similar to the effect of a competing species (Liu *et al*. 2023). The mean of drive loss rate per chasing generation is larger than 0, and there is almost no difference when embryo resistance is between 0 and 0.1 (**Figure 5B**), but there is a substantial decrease when the embryo resistance is larger than 0.1. This shows that limiting the number of r2 resistance alleles does not reduce the drive loss rate when chasing, but this could still be an important way to reduce the chance of chasing in the first place.

We examined the effect of model type, including those based on offspring viability versus adult fecundity, discrete generations versus overlapping generations, the effect of multiple mating, and the effect of different density-dependent growth curves. Our conclusions allowed us to determine why population elimination may be more difficult in spatial mosquito-specific models (Champer *et al*. 2022) than for more commonly used discrete-generation models. These variants could be more or less applicable to particular species, underscoring the critical importance of considering ecology for population suppression systems of gene drives. For our fecundity models, females in favorable low-density positions at the edge of a chasing cluster will disperse offspring equally into all directions, including back in the chasing cluster. In the viability model, offspring will be similarly dispersed, but those further away will have greater survival, thus allow wild-type to advance more quickly into empty areas while preventing the drive from similarly advancing quickly into wild-type. This promotes chasing. Overlapping generations with remating increases effective dispersal of individual alleles due to remating, and previous studies have shown that higher dispersal reduces chasing (Champer *et al*. 2021a). When remating is removed, models with overlapping generations had the same suppression rate as discrete generation models during chasing, though population sizes still tended to be a little higher.

While our model provides a basic understanding of chasing and how various factors can affect it dynamics, real-world populations may have additional factors that we did not consider that can substantially change these dynamics. For example, non-random migration may greatly influence dynamics in chasing situations where organisms are actively responding to spatial variation in population density. Our analysis also used simplified models that had qualitative differences in life history traits, but did not directly represent any particular species such as mosquitoes, nor did we consider spatial variation in movement or carrying capacity (both common in real-world locations). Finally, we did not comprehensively analyze all drive performance parameters, and other types of suppression drives were not considered, which can have large performance differences (Champer *et al*. 2021a; Faber *et al*. 2021; Liu & Champer 2022; Faber *et al*. 2023).

In summary, this study provides a method to quantify chasing population dynamics and explore the effect of drive performance, density dependence, and life history traits. Here, we used a female fertility homing drive to research chasing dynamics, but our detection and evaluation system can apply to many other population suppression gene drive systems that have different mechanisms. This study further expands our knowledge on the relationship between spatial population dynamics and gene drive performance, which could be incorporated into more detailed economic or disease models. These considerations will be important for determining the necessary drive performance for success in different species, and whether any future suppression gene drives could potentially be considered as release candidates.

## Acknowledgements

The cluster-based data collection was assisted by High-Performance Computing Platform of the Center for Life Science at Peking University. This study was supported by the Center for Life Sciences and the National Science Foundation of China (grant 32270672). PWM was supported by the National Institutes of Health under award R35GM152242.

## Supplemental Information

**Table S1.**
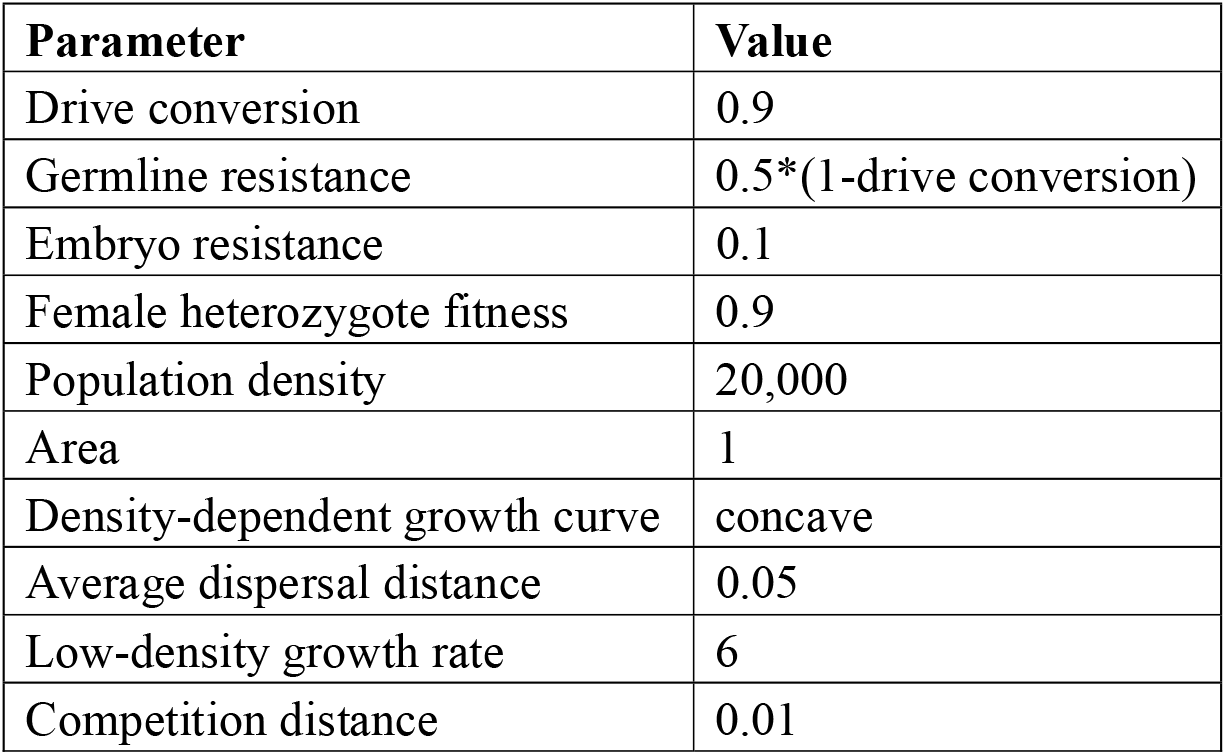
Default parameter values for simulations.

**Figure S1.**
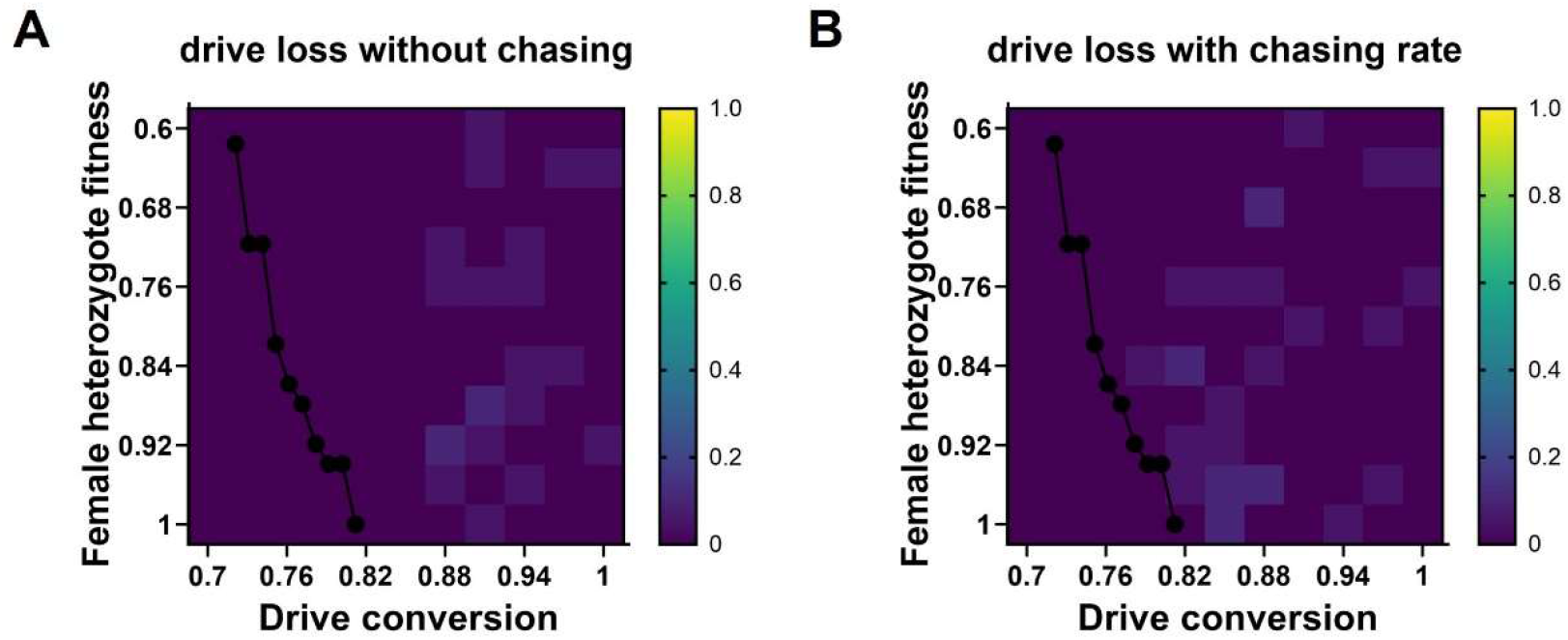
Drive loss rates. Simulations use the discrete-generation fecundity model with 20 simulations for each combination of varying drive conversion and female heterozygote fitness. Other parameters are at their default values. Heatmaps show rates of (**A**) drive loss without chasing and (**B**) drive loss during chasing.

**Figure S2.**
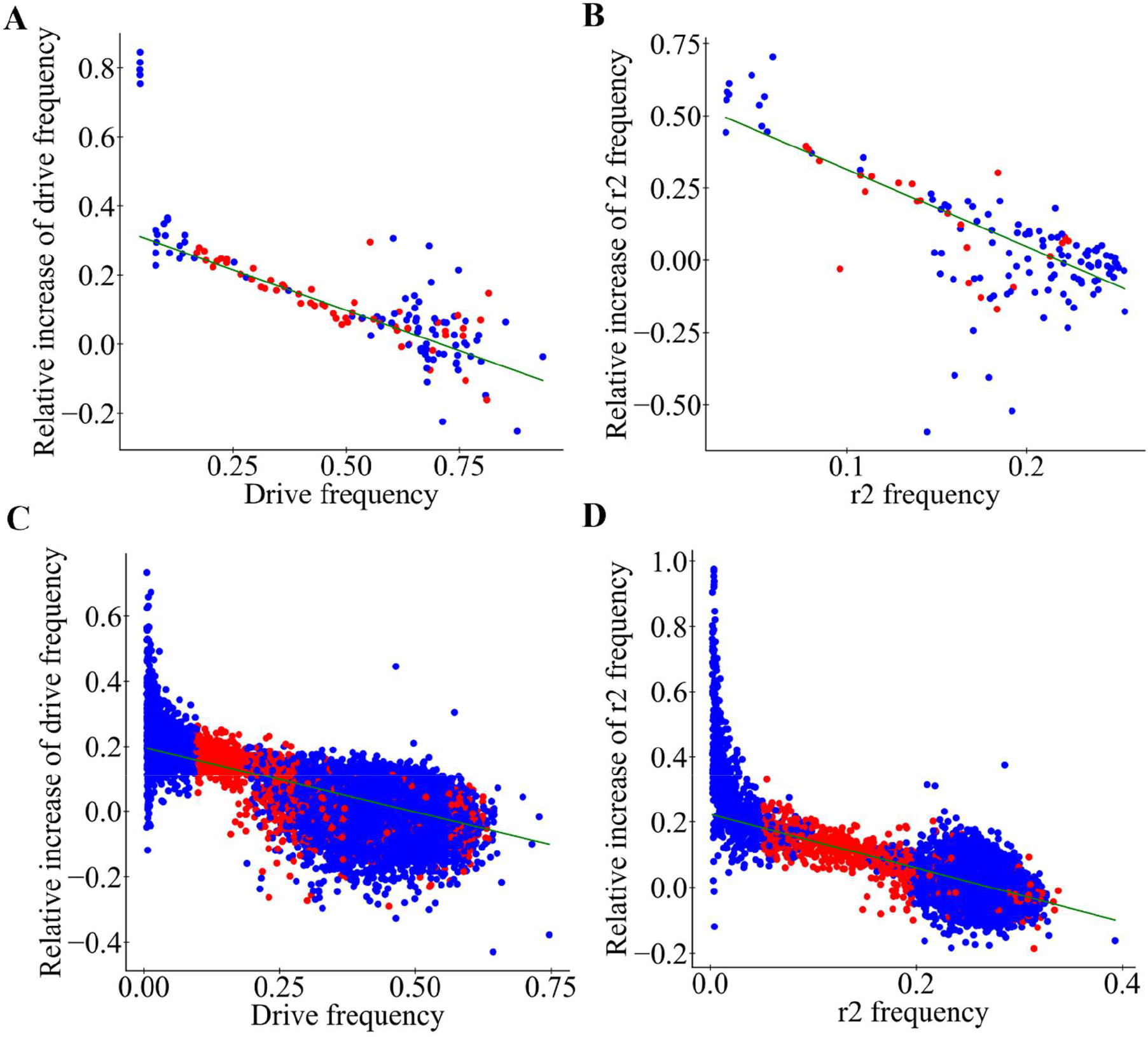
Sample data for drive performance evaluation regression. We released drive into a discrete-generation fecundity population model with a drive conversion of 0.88, embryo resistance of 0.8, and other parameters at default. In a panmictic model, the relationship between (**A**) drive or (**B**) r2 frequency and their relative increase rate is shown. In a spatial model with a central release, the relationship between (**C**) drive or (**D**) r2 frequency and their relative increase rate is shown. Red dots represent chosen samples for linear regression.

**Figure S3.**
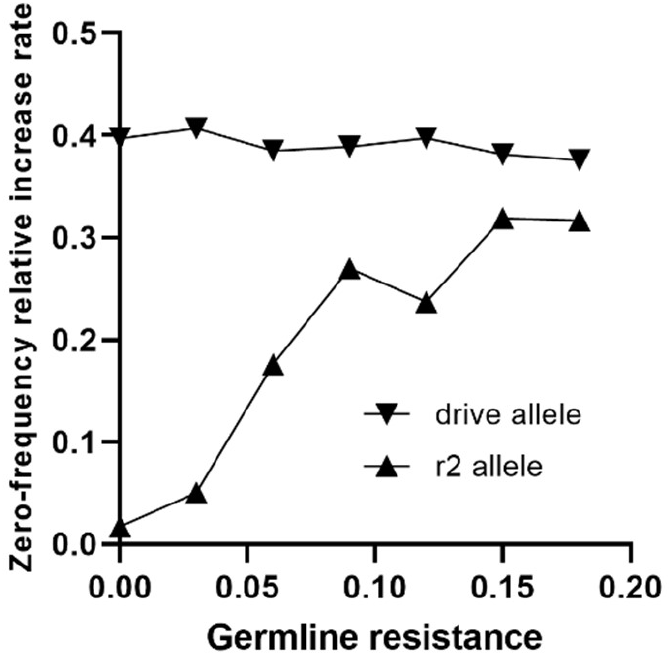
The zero-frequency relative increase rate of drive alleles and resistance alleles. The drive conversion is 0.82, and other parameters are at their default values. The graph shows the relative increase rate when the drive and resistance alleles are at very low frequencies, based on trendlines of drive performance data (see Figure S2).

**Figure S4.**
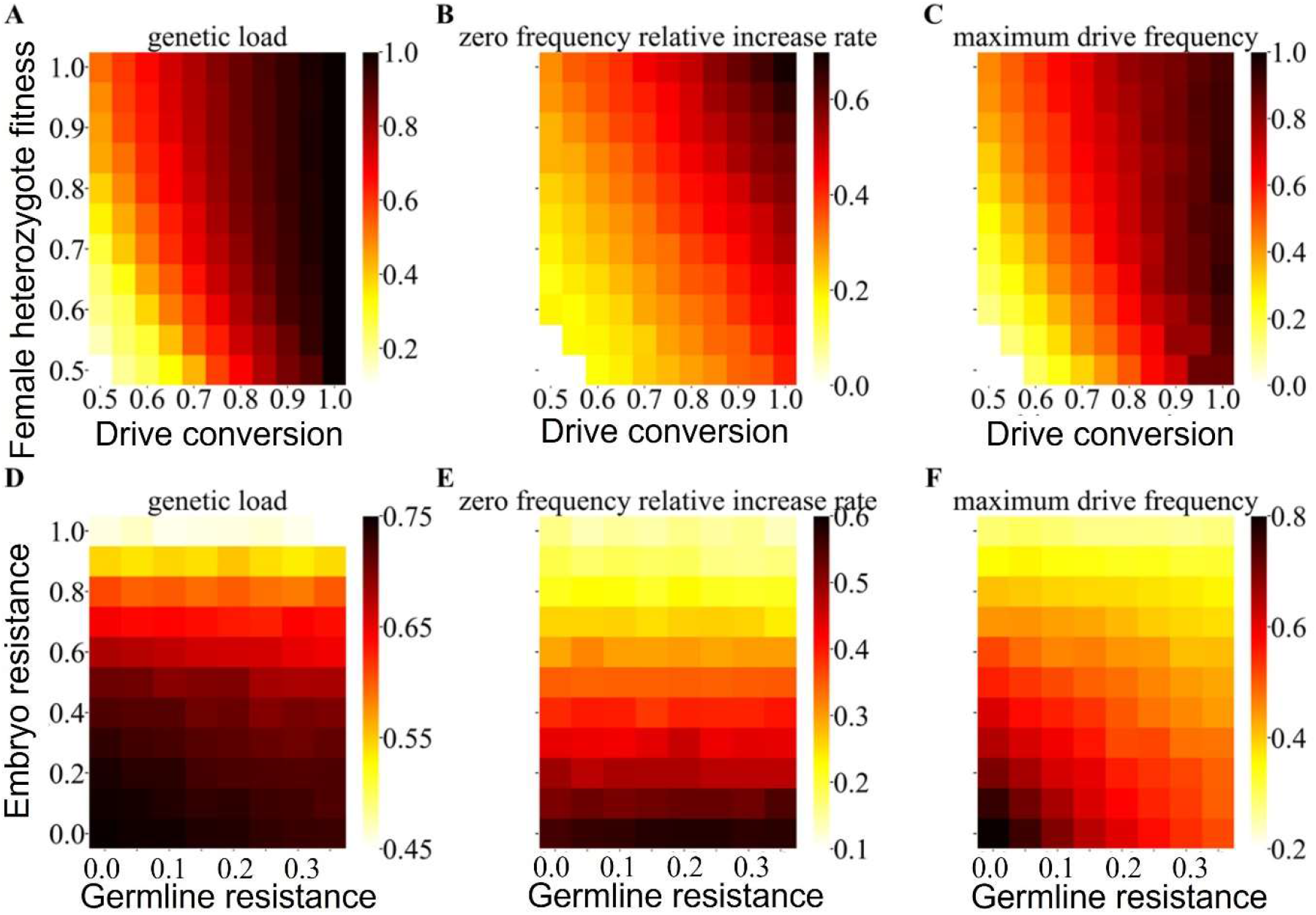
Suppression drive characteristics. With a germline resistance of 0.0 and embryo resistance of 0.4 in the discrete-generation panmictic model, heatmaps show (**A**) the genetic load, (**B**) the zero-frequency relative increase rate of the drive, and (**C**) maximum drive frequency at equilibrium for varying drive conversion and female heterozygote fitness. With a drive conversion of 0.65 and female heterozygote fitness of 0.95, heatmaps show (**D**) the genetic load, (**E**) the zero-frequency relative increase rate of the drive, and (**F**) the maximum drive frequency at equilibrium with varying germline resistance and embryo resistance.

**Figure S5.**
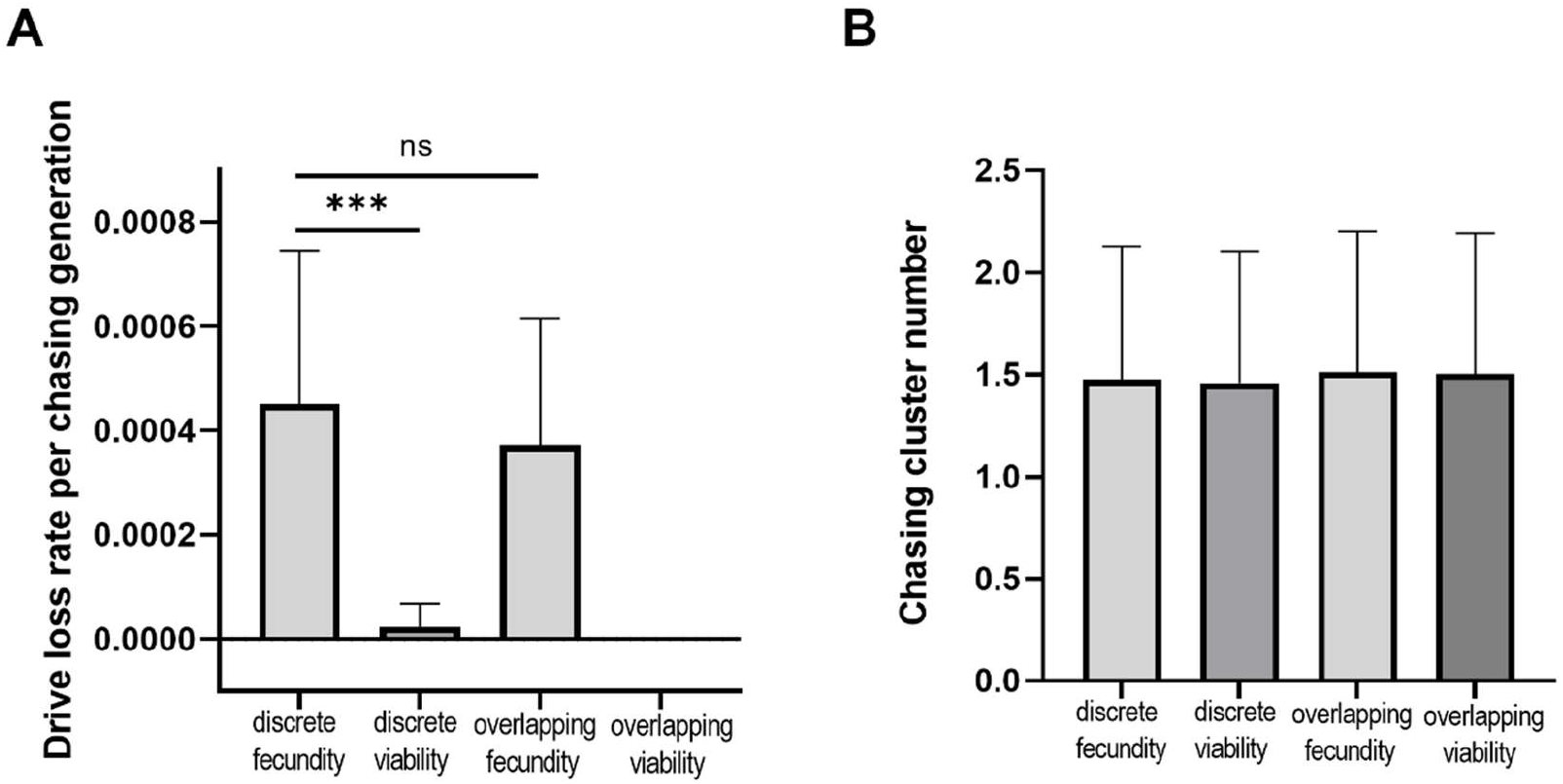
Drive loss rates and number of chasing clusters in different models. With a drive conversion of 0.86, female heterozygote fitness of 0.8, and other parameters at default values, the (**A**) drive loss rates per generation of chasing and the (**B**) average number of chasing clusters are displayed. Error bars in (**A**) are 95% confidence intervals of the estimated success probability from the Bernoulli distribution. Error bars in (**B**) represent the standard deviation. Chi-squared test indicates are ns: not significant, * *p* < 0.05, ** *p* < 0.01, *** *p* < 0.001.

**Figure S6.**
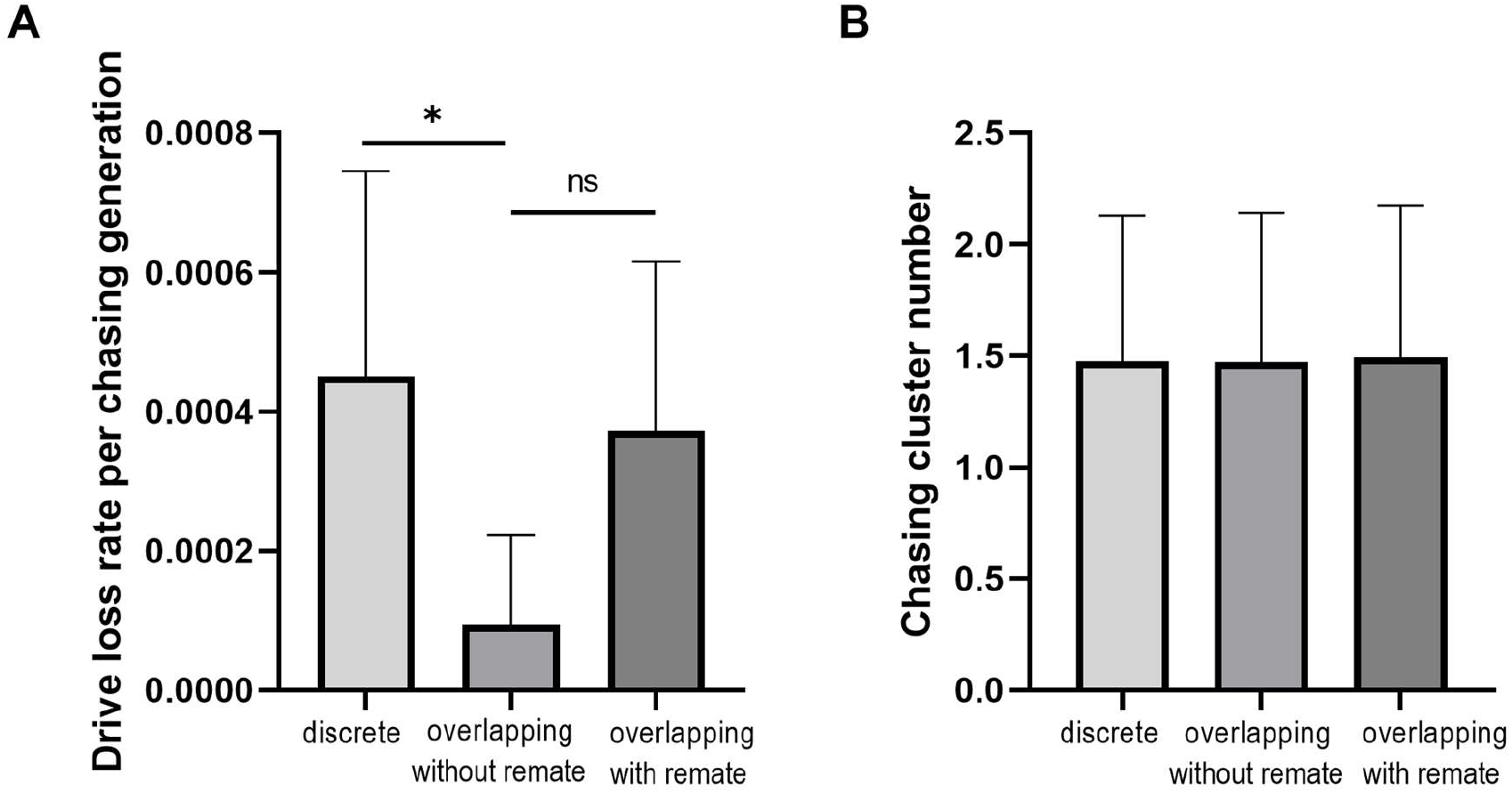
The influence of remating on the drive loss rate during chasing and number of chasing clusters. With a drive conversion of 0.86, female heterozygote fitness of 0.8, and other parameters at their default values, the (**A**) drive loss rates per generation of chasing and the (**B**) average number of chasing clusters are displayed. Error bars in (**A**) are 95% confidence intervals of the estimated success probability from the Bernoulli distribution. Error bars in (**B**) represent the standard deviation. Fisher’s exact test indicates ns: not significant, * *p* < 0.05.

**Figure S7.**
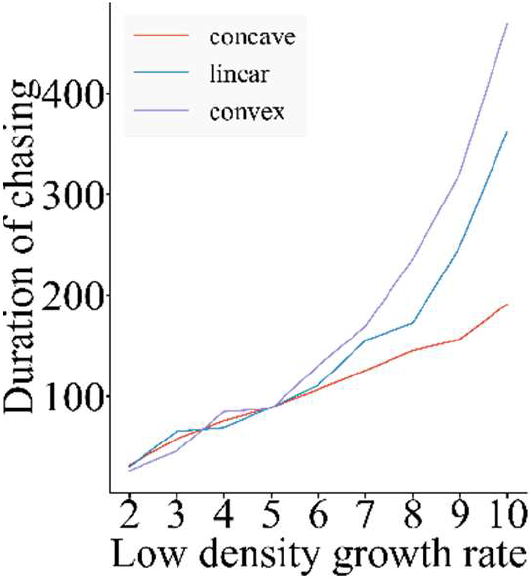
Duration of chasing with varying density growth curve. With a population density of 50,000 and other parameters at their default values, the graph shows the average duration of chasing with variable low-density growth rate and different density-dependent growth curve. Each point is the average of 200 simulations.

